# Multiple motors cooperate to establish and maintain acentrosomal spindle bipolarity in *C. elegans* oocyte meiosis

**DOI:** 10.1101/2021.09.09.459640

**Authors:** Gabriel Cavin-Meza, Michelle M. Kwan, Sarah M. Wignall

## Abstract

While centrosomes organize spindle poles during mitosis, oocyte meiosis can occur in their absence. Spindles in human oocytes frequently fail to maintain bipolarity and consequently undergo chromosome segregation errors, making it important to understand mechanisms that promote acentrosomal spindle stability. To this end, we have optimized the auxin-inducible degron system in *C. elegans* to remove factors from pre-formed oocyte spindles within minutes and assess effects on spindle structure. This approach revealed that dynein is required to maintain the integrity of acentrosomal poles; removal of dynein from bipolar spindles caused pole splaying, and when coupled with a monopolar spindle induced by depletion of kinesin-12 motor KLP-18, dynein depletion led to a complete dissolution of the monopole. Surprisingly, we went on to discover that following monopole disruption, individual chromosomes were able to reorganize local microtubules and re-establish a miniature bipolar spindle that mediated chromosome segregation. This revealed the existence of redundant microtubule sorting forces that are undetectable when KLP-18 and dynein are active. We found that the kinesin-5 family motor BMK-1 provides this force, uncovering the first evidence that kinesin-5 contributes to *C. elegans* meiotic spindle organization. Altogether, our studies have revealed how multiple motors are working synchronously to establish and maintain bipolarity in the absence of centrosomes.

## INTRODUCTION

Centrosomes serve as prominent microtubule organizing centers (MTOCs) during mitosis and provide a clear structural blueprint for the formation of a bipolar spindle. However, oocytes of many species lack centrosomes. In human oocytes, it has been shown that acentrosomal spindles are highly unstable; poles go through an extended period where they can split apart and then come back together, and spindles that display this instability have a high incidence of chromosome segregation errors (Holubcova et al., 2015). The causes of this instability and the mechanisms by which acentrosomal spindles are stabilized in the absence of centrosomes remain poorly understood.

*C. elegans* is an ideal model to investigate these questions, given similarities between worm and human oocytes. Specifically, while acentrosomal spindle assembly in some organisms relies on discrete microtubule organizing centers that help organize spindle poles (Schuh and Ellenberg, 2007), *C. elegans* and human oocytes do not appear to contain these structures (Holubcova et al., 2015; Wolff et al., 2016). In *C. elegans*, microtubules are instead nucleated in proximity to the disassembling nuclear envelope in a spherical cage-like structure. Microtubule minus ends are then pushed outwards to the periphery of the array by the kinesin-12 family motor KLP-18, where they form multiple poles with clearly defined clusters of minus ends. Spindle poles then coalesce, eventually forming a bipolar structure with aligned chromosomes (Wolff et al., 2016; Gigant et al., 2017). Shortly after establishing a metaphase plate, the spindle shortens and rotates perpendicular to the cortex and chromosomes begin to segregate in anaphase (Albertson and Thomson, 1993). The cortical set of homologous chromosomes are discarded as a polar body, and the remaining sister chromatids undergo a second round of meiosis to generate the final set of maternal DNA.

While these stages of acentrosomal spindle assembly have been documented in multiple studies, the mechanisms by which microtubules organize into stable acentrosomal poles in this organism are not well understood. It would stand to reason that mitotic pole proteins are ideal candidates for also promoting meiotic pole stability; many proteins that serve important functions in *C. elegans* mitotic spindle assembly can be found within acentrosomal meiotic spindles (reviewed in (Severson et al., 2016; Mullen et al., 2019)). Various microtubule-associated proteins (MAPs) and microtubule motors have been demonstrated to concentrate at poles and contribute to acentrosomal spindle assembly, such as ASPM-1 (Connolly et al., 2014), KLP-7^MCAK^ (Connolly et al., 2015; Han et al., 2015; Gigant et al., 2017) and MEI-1/2^katanin^ (Srayko et al., 2000; McNally et al., 2014). A study using a fast-acting temperature-sensitive *mei-1* mutant demonstrated that katanin is essential to maintain spindle structure (McNally et al., 2014), but whether any other pole associated-proteins are required for spindle maintenance is not known.

Another factor that has been implicated in pole function is the minus-end directed microtubule motor dynein. Studies in other systems have shown that dynein is required for mitotic spindle pole focusing (reviewed in (Borgal and Wakefield, 2018)) and promoting poleward flux of newly formed microtubule minus ends (Elting et al., 2014; Hueschen et al., 2017). In *C. elegans* embryos, cortically-associated dynein is crucial for the proper positioning of both mitotic and acentrosomal meiotic spindles (Ellefson and McNally, 2009; van der Voet et al., 2009; Ellefson and McNally, 2011; Crowder et al., 2015). Moreover, a number of studies have shown that depletion of dynein or its cofactor dynactin causes spindle defects in oocytes, such as longer spindles with pole defects (Yang et al., 2005; Ellefson and McNally, 2009; Ellefson and McNally, 2011; Crowder et al., 2015; Muscat et al., 2015). However, since dynein is essential for development, these prior studies relied on either partial depletion or on temperature-sensitive mutants, which might not be completely null, so the effect of complete dynein inhibition on spindle assembly has not been assessed. Moreover, whether dynein is required to maintain acentrosomal pole integrity has not been tested. Thus, there is a need for an approach that can completely remove dynein function from oocytes in a conditional manner.

Here, we have addressed this challenge using the auxin-inducible-degron (AID) system (Zhang et al., 2015). We found that this method could efficiently remove dynein from oocytes within minutes, enabling us to remove this protein either prior to spindle assembly or after bipolarity has already been established. This approach revealed that dynein is essential for both the formation and stabilization of acentrosomal spindle poles. Removing dynein from monopolar spindles caused catastrophic breakdown of the monopole, further supporting a role for dynein in pole integrity. Moreover, this experiment also revealed a striking phenotype that led to additional insights into acentrosomal spindle function. Specifically, we found that following disruption of the monopole, individual chromosomes were able to reorganize local microtubule bundles into a bipolar spindle that could mediate anaphase-like segregation. This phenotype provided an opportunity to identify proteins contributing to spindle bipolarity that are masked under normal conditions. Through this assay, we identified kinesin-5 motor BMK-1 as an outward sorting force on microtubules, providing the first evidence that kinesin-5 contributes to spindle assembly in *C. elegans*. Altogether, these studies have furthered our understanding of the motor forces that contribute to acentrosomal spindle assembly and maintenance during meiosis.

## RESULTS

### Dynein is localized to acentrosomal spindles and is required for pole coalescence

Since dynein is required for multiple cellular processes in *C. elegans*, previous studies using RNAi have used partial depletion to avoid impacting worm development (Yang et al., 2005; Ellefson and McNally, 2009; Ellefson and McNally, 2011; Muscat et al., 2015; McNally et al., 2016; Laband et al., 2017). In an attempt to achieve more complete depletion, we utilized a strain in which the heavy chain of dynein, DHC-1, was tagged at the endogenous locus with both GFP and a degron tag. This strain also expressed the ubiquitin ligase TIR1 using the *sun-1* promotor to enable auxin-inducible degradation (AID) of DHC-1 specifically in the germ line (Zhang et al., 2015) (**Figure 1A**; we refer to this strain as “Dynein AID”). With this system, we are able to deplete dynein on a shorter time scale than RNAi, allowing us to assess the effects of full dynein depletion on spindle assembly and stability without affecting prior processes. To validate that this strain worked for efficient DHC-1 depletion, we imaged one-cell stage mitotic embryos using immunofluorescence (IF) and looked for canonical dynein depletion phenotypes, such as failure to separate centrosomes and improper mitotic spindle positioning (Gonczy et al., 1999; Nguyen-Ngoc et al., 2007; Kiyomitsu and Cheeseman, 2013). After a 4-hour incubation of adult Dynein AID worms on auxin-containing plates, we observed clear defects in centrosome separation and spindle positioning along the A-P axis and dynein was undetectable (**Figure 1B**), confirming strong depletion in embryos under conditions where germline and worm development are not affected.

**Figure 1.**
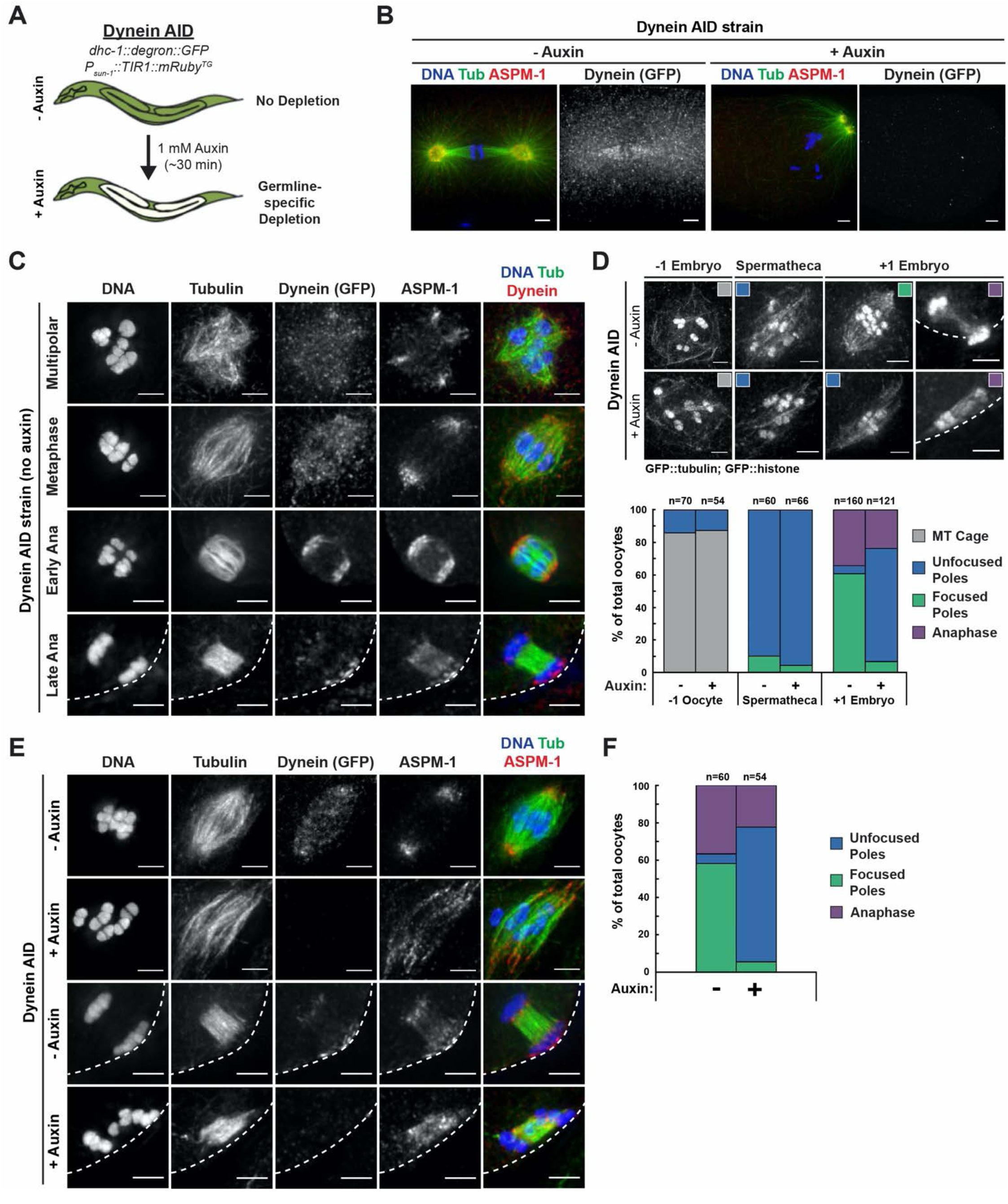
Dynein is required for acentrosomal pole focusing. (A) Schematic representation of the Dynein AID system for DHC-1 depletion in the *C. elegans* germ line. (B) IF imaging of one-cell mitotically-dividing embryos shows that auxin treatment causes efficient dynein depletion and canonical mitotic spindle defects. Shown are tubulin (green), DNA (blue), ASPM-1 (red) and dynein (not shown in merge). (C) IF imaging of oocyte spindles in the Dynein AID strain shows that dynein is localized to the spindle, with increasing enrichment at acentrosomal poles at the anaphase transition; shown are tubulin (green), DNA (blue), dynein (red) and ASPM-1 (not shown in merge). Cortex is represented by dashed line. (D) Representative images of oocyte spindles (GFP::tubulin and GFP::histone) in germline counting and corresponding quantifications; auxin treatment leads to splayed poles and spindle rotation defects. Cortex is represented by dashed line. (E) IF imaging of Dynein AID conditions showing effects of 4-hour auxin treatment on metaphase (second row) and anaphase (bottom row); shown are tubulin (green), DNA (blue), ASPM-1 (red) and dynein (not shown in merge). ASPM-1 labeling supports initial observations of splayed poles seen in germline counting. (F) Quantifications of IF imaging shown in (E); meiotic spindles have significantly splayed poles upon auxin treatment. All scale bars = 2.5µm.

To further validate the Dynein AID strain, we assessed the localization of tagged dynein throughout oocyte meiosis and found that the pattern was consistent with previous studies (**Figure 1C**) (Ellefson and McNally, 2009; van der Voet et al., 2009; Ellefson and McNally, 2011; Crowder et al., 2015). Dynein was weakly associated with forming and bipolar spindles with a slight enrichment at acentrosomal poles labeled by the microtubule minus marker ASPM-1, and then became strongly enriched at poles during early anaphase; this enrichment decreased by late anaphase, with a population being retained near the cortex. Live imaging corroborated our fixed imaging (**Video 1**) and also revealed a population of dynein at kinetochores, as has been shown in other live imaging studies (McNally et al., 2016; Danlasky et al., 2020).

Given this initial validation, we created a Dynein AID strain that also expressed GFP::tubulin and GFP::histone and then assessed the effects of dynein depletion in oocytes by quantifying spindle morphology in intact worms (**Figure 1D**). The *C. elegans* germ line is arranged in a production-line fashion. Oocytes are fertilized in the −1 position of the germ line, and then spindles assemble as the newly fertilized embryo moves through the spermatheca and into the +1 position (Wolff et al., 2016). Under control conditions, bipolar spindles with focused poles were prevalent by the time the embryo reached the +1 position (**Figure 1D**). However, following 4 hours of auxin treatment, spindles mostly had unfocused poles; a large number of spindles displayed microtubule bundles arranged in a relatively parallel array, similar to what was observed following depletion of dynactin component DNC-1 in a previous study (Crowder et al., 2015). Imaging of ASPM-1 confirmed this phenotype; following 4-hour auxin treatment, microtubule minus ends lacked coalesced points between the ends of microtubule bundles (**Figure 1E, 1F**). Moreover, we also saw defects in spindle rotation in anaphase, consistent with previous studies (Ellefson and McNally, 2009; van der Voet et al., 2009; Ellefson and McNally, 2011; Crowder et al., 2015). These findings confirm the prediction that dynein is required for pole focusing during spindle assembly in oocytes, and validate the use of the Dynein AID strain to study dynein in oocytes and embryos.

### Dynein is required throughout meiosis to maintain focused poles

After corroborating that dynein was required to focus acentrosomal poles during spindle assembly, we next sought to determine whether dynein was required to maintain focused poles. We therefore arrested oocytes in Metaphase I to enrich for pre-formed spindles (by depleting anaphase-promoting complex component EMB-30 (Furuta et al., 2000)) and then soaked worms in auxin for 25-30 minutes to acutely deplete dynein. Under these conditions, nearly all spindles had severely splayed poles (**Figure 2A, 2B**); microtubule bundles ran laterally alongside the bivalents, but the ends of the bundles were not brought together into focused poles. While ASPM-1 was enriched at these poles, it also was more dispersed throughout the spindle, suggesting a decrease in poleward transport of microtubule minus ends that has been reported in dynein-depleted mitotic spindles (Hueschen et al., 2017). Moreover, spindles appeared significantly longer after acute dynein depletion (**Figure 2 – figure supplement 1A**). We also observed identical spindle phenotypes in the absence of the metaphase arrest (**Figure 2A, 2B, Figure 2 – figure supplement 1B**).

**Figure 2.**
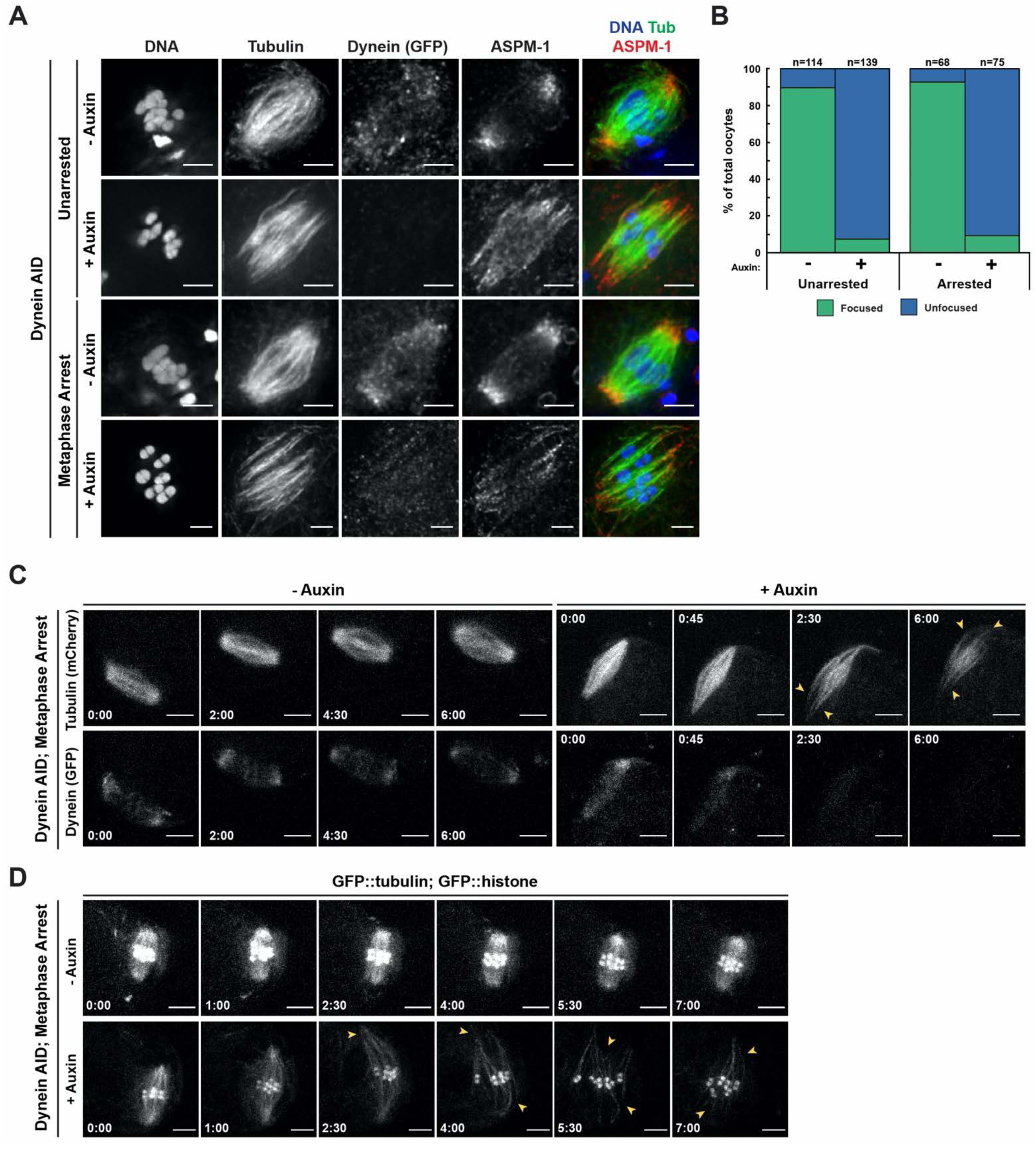
Dynein is required to maintain focused spindle poles. (A) IF imaging of oocyte spindles in control or metaphase-arrest *(emb-30(RNAi))* conditions; shown are tubulin (green), DNA (blue), ASPM-1 (red) and dynein (not shown in merge). Acentrosomal poles become unfocused upon acute dynein depletion. Scale bars = 2.5µm. (B) Quantifications of bipolar spindles imaged in (A) shows that addition of auxin causes nearly all spindle poles to unfocus and splay, both with and without the metaphase arrest. Further quantification of changes in spindle length and microtubule minus end distribution upon auxin treatment can be seen in Figure Supplement 1A and 1B. (C) *ex utero* live imaging of metaphase-arrested spindles; vehicle-treated control oocytes are shown on the left, auxin-treated oocytes are shown on the right. Addition of auxin causes rapid depletion of dynein (labeled with GFP, bottom row) and dynamic splaying of both spindle poles (shown using mCherry::tubulin, top row; arrowheads). Time elapsed shown in (min):(sec). Scale bars = 5µm. (D) *ex utero* live imaging of metaphase-arrested spindles, shown using GFP::tubulin and GFP::histone; addition of auxin (bottom row) causes dynamic splaying of poles (arrowheads) but does not grossly disrupt chromosome alignment; vehicle-treated control oocytes are shown on the top row. Time elapsed shown in (min):(sec). Scale bars = 5µm. Additional *ex utero* imaging of unarrested embryos treated with auxin can be seen in Figure Supplement 2.

To confirm these results and to assess the dynamics of pole splaying, we turned to live *ex utero* imaging of embryos to film acute dynein AID depletion from metaphase-arrested spindles. We generated a Dynein AID strain expressing mCherry::tubulin so that we could visualize the spindle while dynein, tagged with GFP, was depleted. In the absence of auxin, there was no noticeable change in dynein localization or brightness throughout the imaging timeframe (**Figure 2C**, **Video 2**). In contrast, when embryos were dissected into auxin, dynein appeared depleted from meiotic spindles within roughly three minutes and acentrosomal poles began to splay (**Figure 2C, Video 3**). Splaying occurred at both poles (**Figure 2C, arrowheads**) and spindle length increased over time. To visualize the movement and alignment of chromosomes during this process, we repeated this experiment in our GFP::tubulin and GFP::histone Dynein AID strain (**Figure 2D)**. As before, metaphase-arrested spindles were quickly disrupted upon dissection into auxin solution (**Videos 4 and 5**); poles began to splay (**Figure 2D, arrowheads**) and spindles again lengthened. Despite these changes, chromosomes and associated microtubule bundles were able to remain relatively aligned. To verify that the phenotypes were not artifacts of metaphase arrest, we repeated this experiment without *emb-30(RNAi)* and saw the same pole defects (**Figure 2 – figure supplement 2, Videos 6 and 7**). These embryos subsequently progressed into anaphase, albeit with defects in spindle positioning, as observed in fixed imaging. Altogether, these data demonstrate that dynein is not only required to focus poles during meiotic spindle assembly, but is also required to maintain the integrity of acentrosomal poles once they have been established.

### Dynein acts at spindle poles to promote pole formation and integrity

Despite the pole defects observed following dynein depletion, spindle microtubules remained aligned along a single axis and bipolarity was not lost, likely because microtubule bundles in the central region remained intact to stabilize the spindle. To test this hypothesis, we sought to deplete dynein from a monopolar spindle, generated by depleting the kinesin-12 motor KLP-18; these structures have a single focused pole and therefore no area of microtubule overlap that could stabilize the structure if the pole is disrupted (Wignall and Villeneuve, 2009). Previously, we found that partial dynein inhibition on a monopolar spindle led to chromosomes moving a substantial distance away from the center of intact monopoles (Muscat et al., 2015); the AID approach now allows us to revisit these results with more complete dynein depletion.

First, we quantified embryos in intact dynein AID worms treated with *klp-18(RNAi)*, and we found that long-term (4 hour) auxin-treated embryos rarely displayed a monopolar spindle (**Figure 3A**). Instead, chromosomes were dispersed throughout the embryo, retaining association to some microtubule bundles. IF imaging following acute auxin treatment confirmed this result; ASPM-1 labeling revealed that monopoles had begun breaking apart, ejecting chromosomes and associated microtubule bundles into the cytoplasm (**Figure 3B**). In some cases, the entirety of the monopole disappeared and all chromosomes retained their own laterally-associated microtubule bundles. We observed the same phenotype using live *ex utero* imaging; when embryos containing monopolar spindles were dissected into auxin solution, we observed rapid breakdown of the monopole on a timescale similar to that required for full dynein depletion (about three minutes) in previous experiments (**Figure 4A, 4B, and Video 8**). As the monopolar spindle began to fragment, microtubule bundles clearly remained associated with individual chromosomes via lateral associations (**Figure 4A, arrowheads**), as seen in fixed imaging (**Figure 3B**). These results demonstrate an essential role for dynein in maintaining pole integrity and suggest that in the case of dynein depletion from bipolar spindles, the intact overlap zone stabilizes the spindle even though poles are completely disrupted.

**Figure 3.**
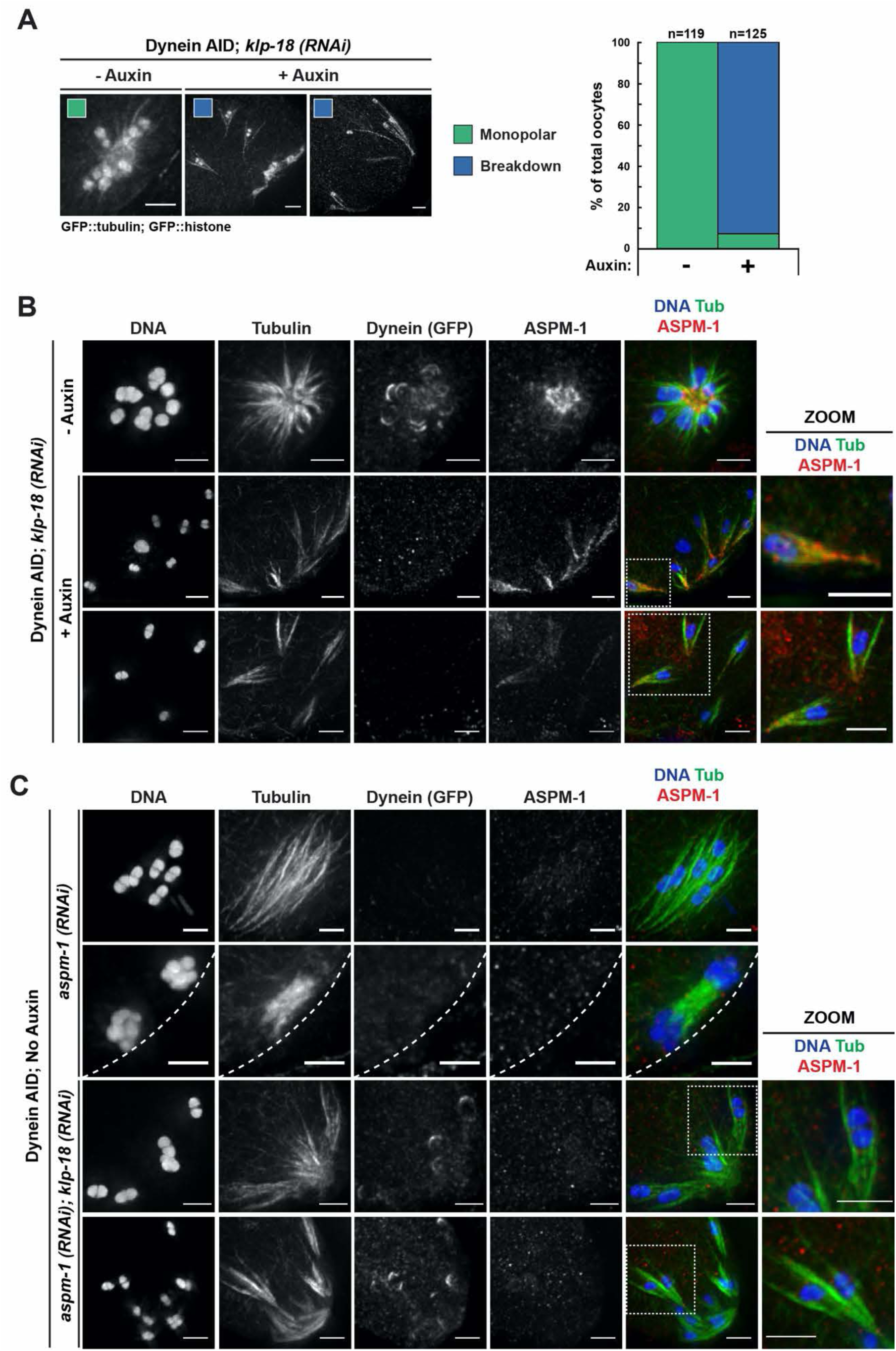
Monopolar spindles breakdown upon dynein depletion. (A) Representative images of *klp-18(RNAi)* oocyte spindles (GFP::tubulin and GFP::histone) in germline counting and corresponding quantifications; dynein depletion leads to dissolution of monopole, releasing chromosomes with associated microtubule bundles into the cytoplasm. (B) IF imaging of monopole breakdown after dynein depletion; shown are tubulin (green), DNA (blue), ASPM-1 (red) and dynein (not shown in merge). Chromosomes released into the cytoplasm retain lateral microtubule associations (zooms). (C) IF imaging of Dynein AID embryos following either *aspm-1* single RNAi or *aspm-1; klp-18* double RNAi; shown are tubulin (green), DNA (blue), ASPM-1 (red) and dynein (not shown in merge). Bipolar spindles exhibit significant splaying of spindle poles and spindle rotation defects in anaphase (top two rows). Double RNAi causes partial breakdown of monopolar spindles (bottom two rows). Cortex represented by dashed line. All scale bars = 2.5μm. Additional conditions involving *lin-5(RNAi)* can be seen in Figure Supplement 1A and 1B.

**Figure 4.**
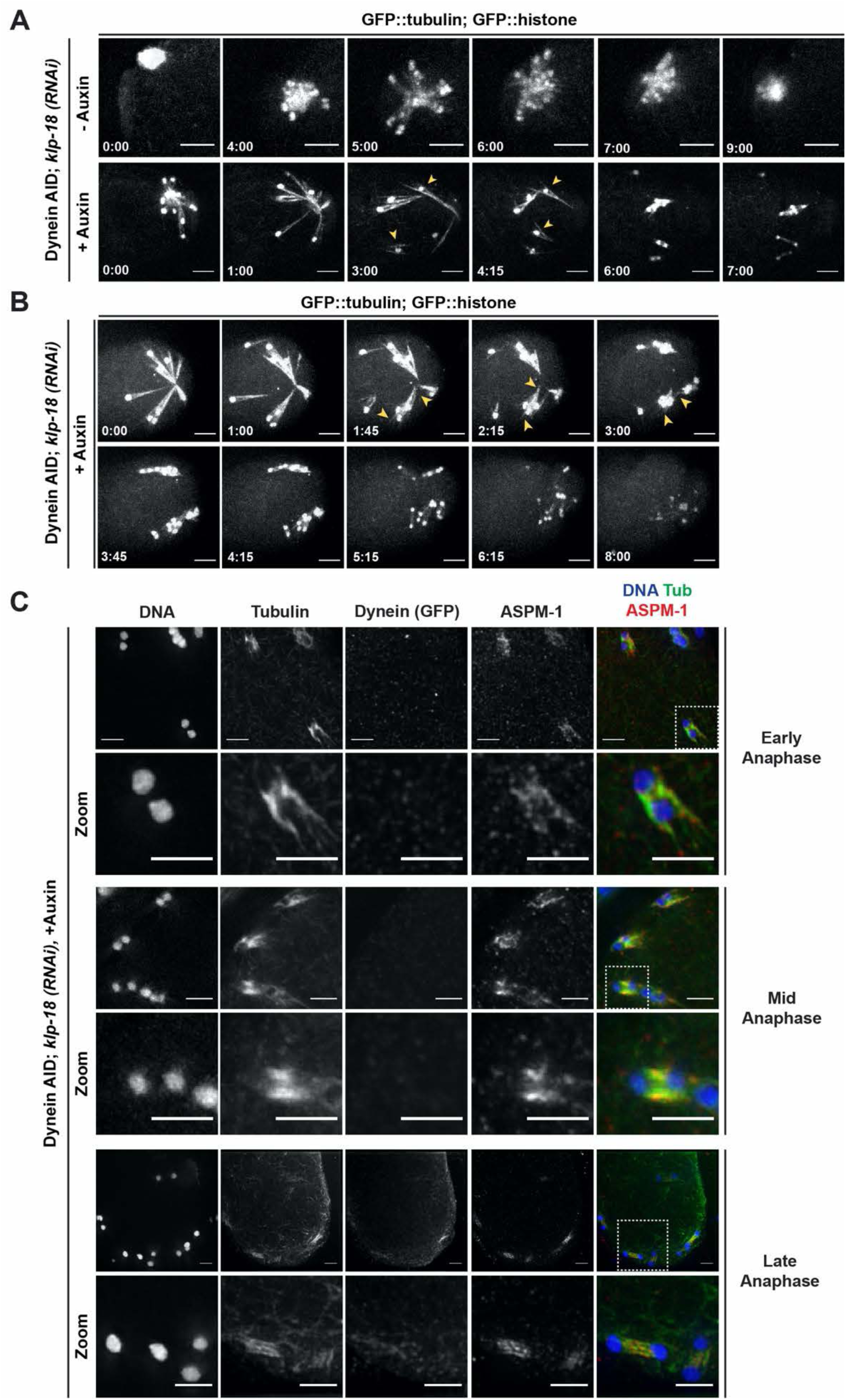
Individualized chromosomes undergo anaphase-like segregation in the absence of KLP-18 and dynein. (A) *ex utero* live imaging of GFP::tubulin and GFP::histone in control and *klp-18(RNAi)* conditions. In the control (top row), a monopolar spindle forms and then chromosomes move back towards the monopole in anaphase, as previously described (Muscat et al., 2015). Addition of auxin to acutely deplete dynein (bottom row) leads to rapid breakdown of monopolar spindles. Microtubule bundles remain laterally associated with chromosomes (arrowheads) prior to anaphase-like segregation. Time elapsed shown in (min):(sec). Scale bars = 5µm. (B) Another example of acute auxin treatment, to remove dynein, in *klp-18(RNAi)* conditions; after breakdown of the monopole and reorganization of microtubules (arrowheads), individual bivalents are able to undergo synchronized anaphase-like segregation. Time elapsed shown in (min):(sec). Scale bars = 5µm. (C) IF imaging of miniature anaphases at multiple chromosome segregation distances, representing various stages of anaphase; shown are tubulin (green), DNA (blue), ASPM-1 (red) and dynein (not shown in merge). Segregation events closely resemble different stages of anaphase A and anaphase B. Scale bars = 2.5µm. Quantifications of miniature anaphase segregation distances and rates can be seen in Figure Supplement 1.

The fact that dynein localizes to poles and depletion has such a dramatic effect on pole integrity suggests that dynein might act at the pole itself to stabilize this structure. To test this hypothesis, we sought to delocalize dynein from poles and assess effects on pole integrity. The microtubule-associated protein NuMA has been shown to direct cytoplasmic dynein to microtubule minus ends and to promote pole focusing during mitosis and pole-directed transport of newly formed minus ends (Elting et al., 2014; Hueschen et al., 2017). Additionally, the *C. elegans* homolog of NuMA, LIN-5, has been shown to target dynein to spindle poles in oocytes to promote spindle rotation (van der Voet et al., 2009). Therefore, we depleted LIN-5 using RNAi to determine if delocalizing dynein from poles would have the same phenotypes as dynein depletion. LIN-5 depletion from bipolar spindles caused pole defects that were identical to 4-hour dynein AID (**Figure 3 – figure supplement 1A);** labeling with ASPM-1 revealed splayed poles and dispersed minus ends throughout the spindle (**Figure 3 – figure supplement 1B**).

Additionally, depletion of LIN-5 from monopolar spindles caused breakdown of monopoles (**Figure 3 – figure supplement 1B**). Dynein AID depletion combined with *lin-5(RNAi)* had similar pole defects as *lin-5(RNAi)* or Dynein depletion alone (**Figure 3 – figure supplement 1B**), supporting the view that dynein and LIN-5 operate in the same pathway (van der Voet et al., 2009). Another protein that acts with LIN-5 to target dynein to poles and promote spindle rotation is ASPM-1. Additionally, depletion of ASPM-1 has been shown to result in longer oocyte spindles with splayed poles (van der Voet et al., 2009; Connolly et al., 2014; Laband et al., 2017) (**Figure 3C**), similar to the phenotype of dynein AID (**Figure 1D-F, Figure 2**), which suggests a role in pole focusing. Consistent with this, we found that ASPM-1 depletion caused the breakdown of monopolar spindles (**Figure 3C**), again suggesting that proper localization of dynein to poles is required for pole integrity. Notably, monopole breakdown was not as severe following *lin-5* or *aspm-1* RNAi as that seen in the acute dynein AID depletions; for *lin-5* and *aspm-*1 RNAi conditions, only ∼20% of monopoles completely broke down (15/73 and 14/61, respectively) and roughly half of embryos displayed partial breakdown (44/73 and 29/61, respectively). This might be caused by incomplete depletion due to the variable efficiency of RNAi, or perhaps because dynein is still present in these embryos and could be partially functional even though it is mislocalized. Regardless, our results suggest that dynein localization to spindle poles is required for the efficient focusing and stabilization of these structures.

### Individualized chromosomes can undergo an anaphase-like segregation in the absence of KLP-18 and dynein

In addition to revealing the dynamics of monopole breakdown, our live imaging of dynein-depleted monopolar spindles revealed an intriguing phenotype. These experiments were not done using metaphase arrest, so in control embryos, chromosomes extended outward from the monopole towards microtubule plus ends, and then moved back towards the monopole during anaphase (**Figure 4A and Video 9**), as has been described previously (Muscat et al., 2015). In auxin-treated embryos, as filming continued past the full breakdown of the monopole, we were surprised to see that each individual chromosome began to segregate (**Figure 4A, 4B, Video 8, 10**). This segregation was incredibly striking, as it occurred synchronously across the embryo. The fact that we observed these anaphase-like movements in the absence of both KLP-18 and dynein suggests that neither motor is absolutely required for chromosome segregation in oocytes.

When we compared the segregation distance of chromosomes from these videos to control anaphases (bipolar spindles with no auxin), we found that chromosomes segregated to a similar final distance on the dynein-depleted *klp-18(RNAi)* mini-anaphase spindles (**Figure 4 – figure supplement 1**). However, mini anaphases did not appear to segregate as far as dynein-depleted bipolar anaphases, where KLP-18 was not depleted. This observation raises the possibility that, even though chromosomes can segregate without KLP-18, this motor may normally contribute to anaphase spindle elongation through microtubule sliding. This contribution has been difficult to test due to the monopolar spindle phenotype that is quickly generated upon removal of KLP-18 (Wolff et al., 2021), which prevents normal anaphase segregation from occurring.

To further characterize these segregations, we performed fixed imaging. This analysis revealed that the mini-anaphases had key hallmarks of normal anaphase spindles; when chromosomes were still close together we observed lateral microtubule bundles running alongside the separating chromosomes (**Figure 4C**, “early anaphase”), and as chromosomes segregated further and spindle length increased, microtubules were largely localized between chromosomes (**Figure 4C**, “late” anaphase). This suggests that, like normal anaphase spindles, these mini spindles undergo morphological changes and anaphase-B-like spindle elongation as they drive chromosomes apart.

### Microtubules can reorganize into mini bipolar spindles in the absence of dynein and KLP-18

The fact that we observed chromosome segregation with hallmarks of normal anaphase led us to hypothesize that some local microtubule reorganization must be occurring after monopole breakdown; without a bipolar distribution of microtubule minus ends, it should not be possible for chromosomes to segregate away from each other in the anaphase-like manner we observed. In live imaging, we were able to visualize a brief window between full monopole breakdown and the initiation of segregation (**Figure 4A, 4B, Video 8, 10**). During this window, microtubule bundles shortened around chromosomes, and microtubule ends could be clearly identified on both sides of individual chromosomes prior to the initial segregation (**Figure 4B, arrowheads**). Subsequently, an anaphase-like spindle could be observed until the completion of segregation. In order to better characterize this possible reorganization, we returned to IF imaging. In a few occasions we were able to capture this transitional period; ASPM-1 labeling could be clearly seen on both sides of these mini spindles, with some enrichment towards areas resembling spindle poles (**Figure 5A**).

**Figure 5.**
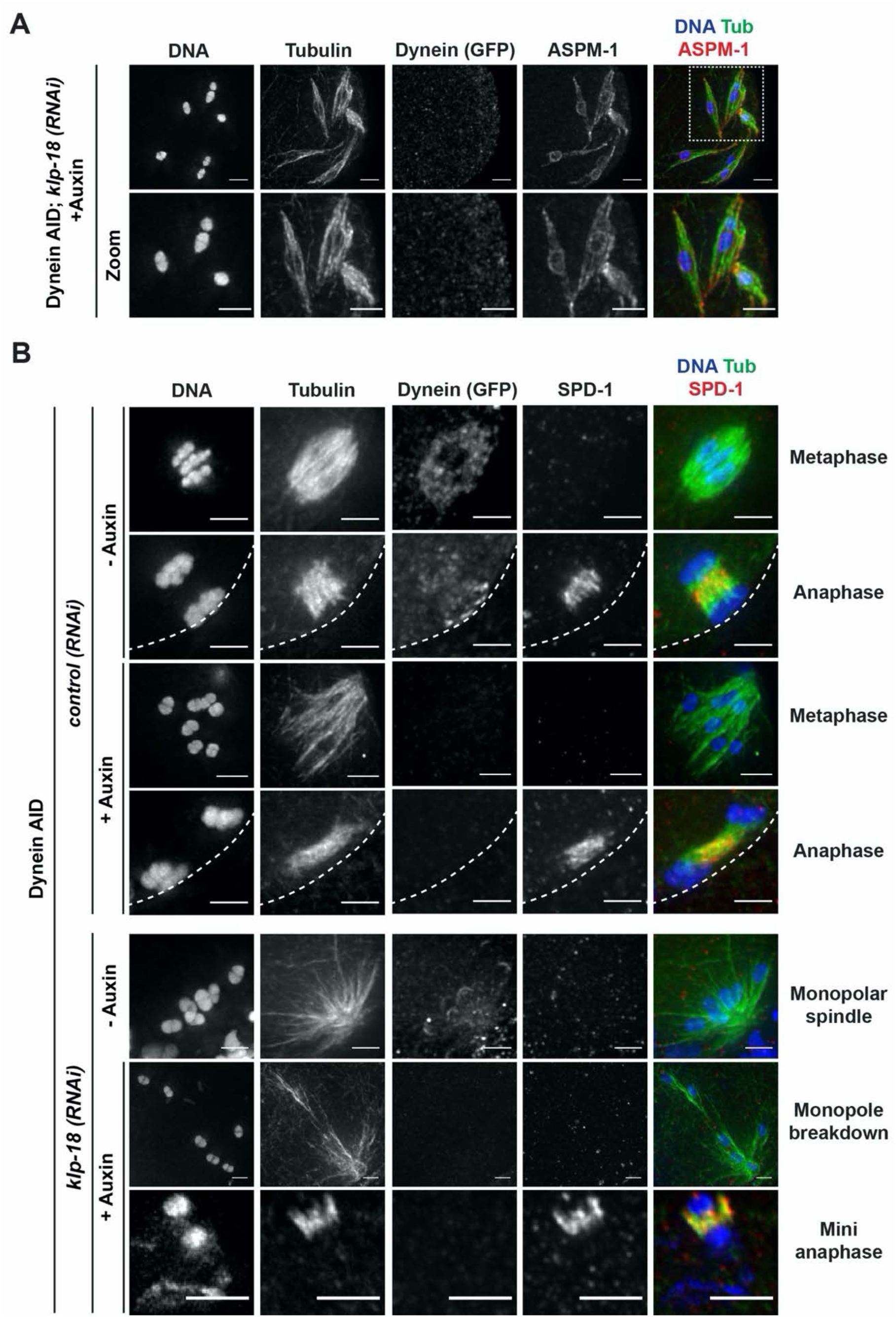
Microtubules reorganize into miniature bipolar spindles that can segregate chromosomes. (A) IF imaging of microtubules (green), DNA (blue), ASPM-1 (red) and dynein (not shown in merge) in monopolar spindle breakdown conditions (*klp-18(RNAi)* + acute dynein AID). Microtubule bundles appear to reorganize around individual chromosomes, seen through ASPM-1 flanking either side of the chromosome; note that in these images ASPM-1 also appears to be on chromosomes, but that is background staining that sometimes occurs with this antibody (Wignall and Villeneuve, 2009) and is not real signal. (B) IF imaging of SPD-1 localization in the Dynein AID strain in control RNAi conditions (rows 1-4) or following *klp-18(RNAi)* (rows 5-7); shown are tubulin (green), DNA (blue), SPD-1 (red) and dynein (not shown in merge). SPD-1 does not localize to spindles in metaphase (rows 1, 3), but localizes to overlapping microtubules in anaphase spindles in the presence or absence of dynein (rows 2, 4). SPD-1 is not localized to monopolar spindles either before or after monopole breakdown (rows 5, 6), but can clearly be seen localized to miniature anaphases (row 7). Cortex is represented by dashed line. All scale bars = 2.5µm. Additional observations of SUMO localization can be seen in Figure Supplement 1.

This imaging suggested that microtubules might reorganize into a mini-bipolar spindle that could facilitate chromosome segregation. To test this idea, we assessed the localization of a well-studied spindle protein, the PRC1 homolog SPD-1. SPD-1 localizes to the anaphase spindle midzone in oocytes, marking a region of antiparallel microtubule overlap (Hattersley et al., 2016; Gigant et al., 2017; Mullen and Wignall, 2017), and we found that SPD-1 was also at this location in the absence of dynein (**Figure 5B,** row 4). In intact and disassembling monopoles, SPD-1 is not localized to microtubules, as it only associates with the spindle in anaphase. However, once chromosomes had been ejected into the cytoplasm and microtubules reorganized around chromosomes, SPD-1 staining could be clearly observed on mini anaphase spindles (**Figure 5B,** bottom row), demonstrating that they have a region of anti-parallel overlap in the center.

In addition to SPD-1, we also assessed the localization of the ring complex (RC), a collection of proteins that forms a ring around the center of each chromosome in oocytes. RCs aid in chromosome congression (Wignall and Villeneuve, 2009; Muscat et al., 2015; Pelisch et al., 2017; Hollis et al., 2020), and then are released from chromosomes during anaphase; they remain as a ring between segregating chromosomes and then are disassembled as anaphase progresses (Dumont et al., 2010; Muscat et al., 2015; Davis-Roca et al., 2017; Pelisch et al., 2017; Davis-Roca et al., 2018; Pelisch et al., 2019). To assess RC behavior, we imaged SUMO, a post-translational modification found in the RC (Pelisch et al., 2017). In control and dynein depletion conditions, SUMO-marked RCs were removed from chromosomes during anaphase and were localized between segregating chromosomes, as expected (**Figure 5 – figure supplement 1**). When introducing monopolar breakdown, SUMO could be seen on the RC as individual chromosomes detached from the dissolving monopole. In mini anaphases, SUMO labeling persisted on what appeared to be a singular disassembling RC between each segregating chromosome pair. Taken together, these findings support the idea that microtubules reorganize into miniature spindles that recapitulate key aspects of normal anaphase.

### The kinesin-5 family motor BMK-1 provides an outward sorting force that allows spindle reorganization in the absence of KLP-18 and dynein

In our monopole breakdown assay, microtubules were able to reorganize around dispersed chromosomes and establish a region of anti-parallel overlap at the center of each mini spindle, suggesting the presence of an activity that can sort microtubules and re-establish bipolarity in the absence of KLP-18 and dynein. We saw this condition as a unique opportunity to probe for redundant motor forces that may be present in a normal meiotic spindle, but would otherwise be masked by the presence of the forces provided by KLP-18 and dynein. One strong candidate was the sole kinesin-5 family motor in *C. elegans*, BMK-1 (Bishop et al., 2005). Kinesin-5 motors have essential roles in establishing spindle bipolarity in many organisms (reviewed in (Mann and Wadsworth, 2019)), but loss of BMK-1 has not been reported to have major phenotypes in *C. elegans;* BMK-1 depletion has no effects on spindle morphology in either oocytes or embryos (Bishop et al., 2005) and only minor effects on chromosome segregation rates (Saunders et al., 2007; Laband et al., 2017). We therefore hypothesized that BMK-1 could be providing a supplementary outward sorting force that is normally masked by the contributions of KLP-18.

To probe this hypothesis, we first sought to confirm the localization of BMK-1 on meiotic spindles and to test if this motor was localized to microtubules under the monopole breakdown conditions. BMK-1 was broadly associated with bipolar metaphase and anaphase spindles, in both control and dynein-depletion conditions (**Figure 6A**). In intact monopolar spindles, BMK-1 also localized to microtubules. Importantly, when we depleted dynein and induced monopole breakdown, BMK-1 was still localized to microtubules (**Figure 6A,** zooms), placing it in a location where it could contribute to microtubule reorganization.

**Figure 6.**
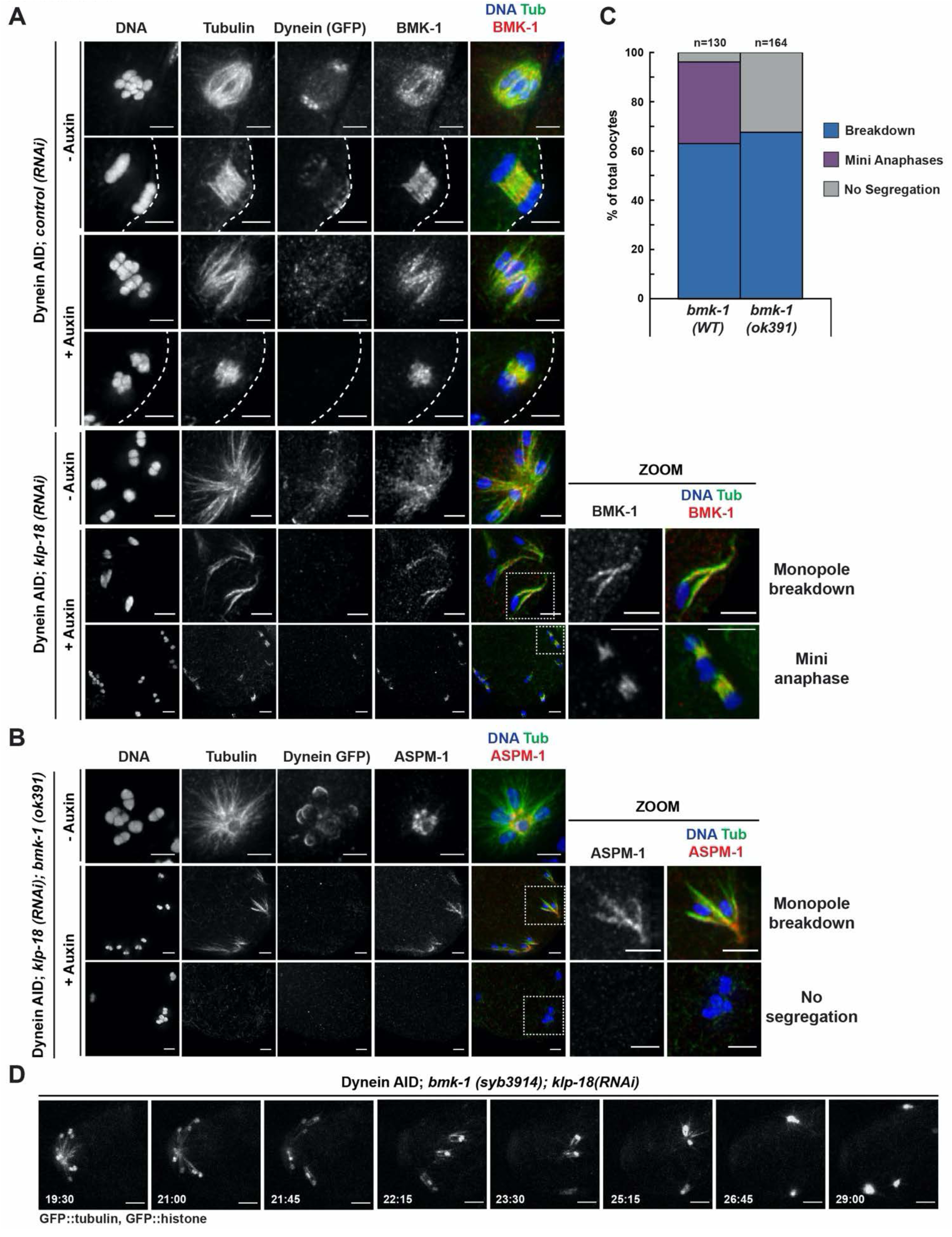
BMK-1 localizes to the meiotic spindle and is required for microtubule reorganization and the formation of miniature anaphases. (A) IF imaging of embryos in either control or *klp-18(RNAi)* conditions in the Dynein AID strain in the presence and absence of acute auxin treatment; shown are tubulin (green), DNA (blue), BMK-1 (red) and dynein (not shown in merge). BMK-1 is localized to spindle microtubules in all conditions, including following monopolar spindle breakdown and in mini anaphases (zoomed images). Cortex is represented by dashed line. Scale bars = 2.5µm. (B) IF imaging of embryos following *klp-18(RNAi)* in the Dynein AID strain lacking functional BMK-1 (*bmk-1(ok391))*; shown are tubulin (green), DNA (blue), ASPM-1 (red) and dynein (not shown in merge). Following monopolar spindle breakdown in the presence of auxin (rows 2, 3), embryos do not contain miniature anaphases and lack chromosome segregation. Scale bars = 2.5µm. (C) Quantifications of images shown in (B) compared to wild type (WT) embryos; monopolar spindles still break down but no miniature anaphases are observed in embryos lacking BMK-1 function. (D) *ex utero* live imaging of GFP::tubulin and GFP::histone following acute auxin treatment to remove dynein in *klp-18(RNAi); bmk-1(syb3914)* conditions; miniature anaphases do not form in the absence of BMK-1. Time elapsed shown in (min):(sec). Scale bars = 5µm. Validation of *bmk-1* mutants via IF imaging can be seen in Figure Supplement 1.

To test if BMK-1 was necessary for the formation of miniature anaphases, we utilized two *bmk-1* mutants: 1) a previously-characterized allele, *bmk-1(ok391)*, that introduces a premature stop codon in the motor domain (Bishop et al., 2005), and 2) a new deletion of the entire *bmk-1* locus generated using CRISPR-Cas9 (*bmk-1(syb3914))*. To validate these deletions, we utilized IF imaging with an α-BMK-1 antibody and confirmed that BMK-1 was no longer present on the meiotic spindle in either mutant (**Figure 6 – figure supplement 1A, 1B**). We then generated monopolar spindles, performed acute dynein AID, and performed IF in the *bmk-1(ok391)* background; in this *bmk-1* mutant we were unable to observe any mini anaphase spindles, even though monopole breakdown still occurred (**Figure 6B, 6C)**; this suggested that removal of BMK-1 function abolished microtubule reorganization and prevented chromosome segregation. To confirm this result, we crossed the *bmk-1* CRISPR deletion mutant into our Dynein AID live imaging strain (expressing GFP::tubulin and GFP::histone to visualize the spindle) and performed *ex utero* imaging to determine if mini anaphases could still form (**Figure 6D**). In *klp-18(RNAi), bmk-1(syb3914)* worms in the absence of auxin, monopolar spindles formed and then chromosomes moved back towards the pole in a manner indistinguishable from normal monopolar anaphase (**Video 11**). When auxin was added to deplete dynein, monopolar spindles broke down as expected and individual chromosomes remained associated with microtubule bundles. However, as time elapsed, there were no signs of segregation, microtubule density decreased, and chromosomes remained in the cytoplasm with no discernable anaphase spindle forming (**Video 12**). These data support the hypothesis that BMK-1 provides outward sorting force on microtubules during oocyte meiosis, redundant to the forces produced by KLP-18.

## DISCUSSION

### Dynein is essential for acentrosomal pole focusing throughout meiosis

Collectively, these data have contributed to a more complete model for how acentrosomal spindles are assembled and stabilized during oocyte meiosis (**Figure 7**). Dynein activity is required throughout the meiotic divisions to establish and maintain focused poles, and its removal via acute AID generated spindle defects within minutes. Dynein depletion increased the length of bipolar oocyte spindles and led to increased dispersion of microtubule minus ends across the spindle. When performing these depletions on a monopolar spindle, the monopole completely broke apart and microtubule bundles were ejected into the cytoplasm, demonstrating that dynein is required to stitch minus ends together into a pole structure. Remarkably, we also found that after the monopole disassembled, microtubules were able to reorganize into a “miniature” spindle capable of facilitating an anaphase-like chromosome segregation. This provides additional evidence that dynein is not essential for chromosome segregation, as has been suggested by previous studies that used partial dynein depletion and/or temperature-sensitive mutants (Muscat et al., 2015; McNally et al., 2016; Laband et al., 2017). In our dynein AID depletions, unarrested bipolar spindles progress through anaphase and mini spindles also undergo anaphase-like segregation, reaching comparable segregation distances to that of normal anaphase spindles. This corroborates a recent study that also used auxin-inducible degradation to assess chromosome segregation (Danlasky et al., 2020); those dynein depletion experiments demonstrate the same segregation distance trends shown here. Moreover, our studies also show that anaphase-B-like spindle elongation can occur in the absence of KLP-18. Although it is possible that KLP-18 may contribute to outward sliding of microtubules during wild type anaphase, our work shows that this motor is not absolutely required for anaphase spindle elongation and demonstrates that there must be other factors that can perform this function.

**Figure 7.**
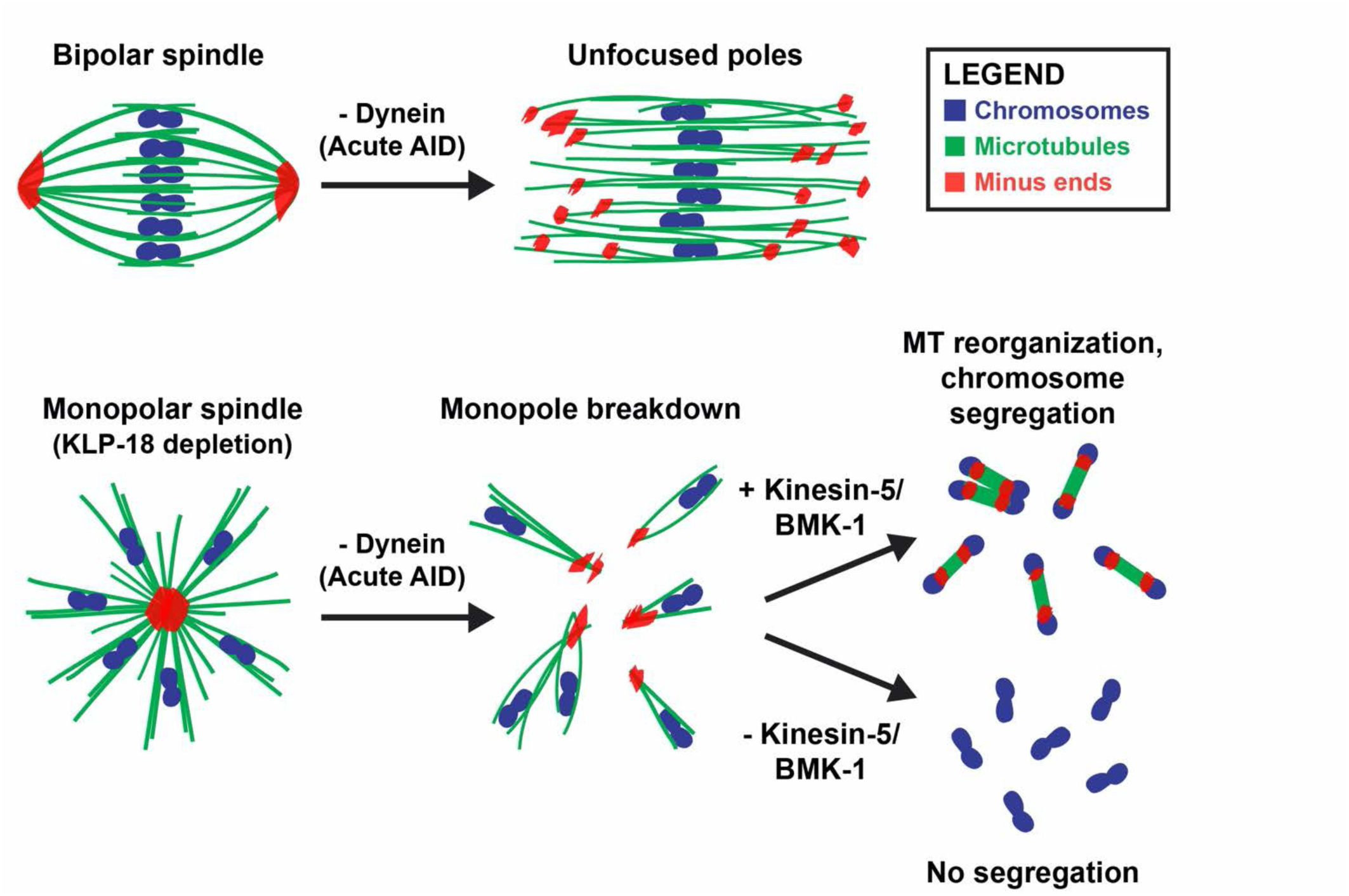
Dynein, KLP-18, and BMK-1 work in concert to establish and maintain spindle bipolarity in *C. elegans* oocyte meiosis. Chromosomes (blue), microtubules (green), and microtubule minus ends (red). Dynein is required throughout the meiotic divisions to maintain focused acentrosomal poles. If removed from stable bipolar spindles (top), poles rapidly unfocus and splay, leading to the same phenotype as long-term depletions. Dynein depletion from monopolar spindles (bottom) ejects individual chromosomes and associated microtubule bundles into the cytoplasm. BMK-1 is able to provide an outward sorting force in the absence of KLP-18 and dynein, enabling reorganization of microtubules into a miniature anaphase spindle, promoting chromosome segregation. In the absence of BMK-1, these miniature anaphases cannot form and anaphase-like segregations no longer occur.

### Multiple motors cooperate in C. elegans oocyte meiosis to effectively form and stabilize an acentrosomal bipolar spindle

Our work also accentuates the importance of balanced forces within a bipolar meiotic spindle; we found that loss of dynein activity had strong phenotypes that manifested rapidly, within two minutes of auxin treatment. Notably, acute dynein depletion caused spindle lengthening, suggesting that dynein may normally provide an inward force on the spindle, as has been demonstrated in previous studies of mitosis (Ferenz et al., 2009; van Heesbeen et al., 2014). Recently, our lab also demonstrated that inactivation of KLP-18 in stable, bipolar spindles caused spindle shortening followed by collapse of microtubule minus ends into a monopolar spindle (Wolff et al., 2021). This collapse is due to a loss of the outward force provided by KLP-18, theoretically enabling the inward force provided by dynein to dominate and highlighting how quickly an imbalance of these motor forces can lead to gross defects in spindle bipolarity. In the current study, removing dynein from oocytes that already lacked KLP-18 resulted in a catastrophic breakdown of the spindle, releasing individual chromosomes into the cytoplasm. Going forward, it would be interesting to acutely remove these proteins at the same time to see if oocyte spindles are able to maintain bipolarity and directly test whether these motors antagonize each other in *C. elegans*.

While KLP-18/kinesin-12 provides the major outward force in *C. elegans* oocyte meiosis, this critical function is performed by kinesin-5 in mouse oocytes (Schuh and Ellenberg, 2007) and in mitosis in many organisms (reviewed in (Mann and Wadsworth, 2019)). Previous studies have shown that dynein and kinesin-5 antagonize each other in these organisms; inhibition of both motors simultaneously enables bipolar spindle formation (Mitchison et al., 2005; Tanenbaum et al., 2008; Ferenz et al., 2009; van Heesbeen et al., 2014). Interestingly, it has been shown that Kif15/kinesin-12 is capable of providing a supplemental outward force that can support bipolarity when kinesin-5 is inhibited (Tanenbaum et al., 2009; Vanneste et al., 2009; Raaijmakers et al., 2012; Sturgill and Ohi, 2013; Sturgill et al., 2016). Here, we have provided evidence that these roles have been reversed in *C. elegans* oocyte meiosis; BMK-1/kinesin-5 appears to be providing a redundant outward sorting force that was only detectable once we had removed KLP-18 and dynein from the meiotic spindle. In the future, it would be fascinating to perform KLP-18/DHC-1 double depletions in a strain lacking BMK-1 function to further probe the relationship between these three motors during acentrosomal spindle formation in *C. elegans*.

While this work has expanded our understanding of the role BMK-1 plays in *C. elegans* oocyte meiosis, further experimentation will be valuable for understanding the exact mechanism of BMK-1 function in the context of a normal, bipolar meiotic spindle. Biophysical assays to determine motor walking speed and force generation would help frame how much BMK-1 contributes in comparison to other meiotic motors such as KLP-18. Also, since kinesin-5 activity is known to be regulated in other systems by kinases and protein-protein interactions (reviewed in (Mann and Wadsworth, 2019)), it would be beneficial to determine if BMK-1 has any interacting partners that provide some regulation of function. Aurora B kinase (AIR-2) has been shown to be required for BMK-1 spindle localization and AIR-2 can phosphorylate BMK-1 *in vitro* (Bishop et al., 2005) implicating AIR-2 in BMK-1 regulation. Inhibition of AIR-2’s kinase activity also leads to collapsed oocyte spindles (Divekar et al., 2021a), consistent with a role for AIR-2 in regulating spindle force generation. However, it is likely that other factors also regulate BMK-1. Finally, a broader understanding of why the relationship of kinesin-5 and kinesin-12 in oocyte meiosis has been seemingly switched between different organisms could provide some valuable context to the optimization of acentrosomal spindles over evolutionary time.

### Different acentrosomal pole proteins have distinct roles in pole coalescence and stability

Our analysis of ASPM-1 and LIN-5 provides further evidence that these proteins are directly influencing acentrosomal pole organization and stability by directing cytoplasmic dynein localization. In either *aspm-1* or *lin-5* RNAi, spindle phenotypes were nearly identical to those seen in Dynein AID experimentation. Additionally, when introducing double depletions with *klp-18(RNAi)*, partial breakdown of monopolar spindles could be observed. Neither double depletion condition was as extreme as Dynein AID phenotypes, but this could be attributed to either inconsistent RNAi efficiency (more common with double depletions) or to partial dynein activity despite it being unable to properly localize to poles. The shared phenotype across depletions of these proteins suggests that these pole proteins function in a single pathway; this provides some credence to the concept that acentrosomal poles could contain a cross-linked network of interacting pole proteins that provide cohesion and stability. However, recent unpublished work from our lab has made it clear that not all pole protein depletions yield the same defects in spindle bipolarity. Acute depletion of ZYG-9, a homolog of the microtubule polymerase XMAP215, disrupts spindle integrity in a manner that does not resemble dynein AID depletions. While dynein depletion splays spindle poles, microtubule bundles appear to remain stable and unperturbed, generating a rectangular spindle that can still progress through meiosis (albeit with spindle rotation defects). However, acute ZYG-9 depletion leads to a disorganized multipolar spindle that is highly prone to segregation errors, suggesting a broader role of ZYG-9 not just restricted to poles.

From these recent studies, it is clear that multiple groups of proteins are required to establish and maintain acentrosomal poles. However, despite having similar or nearly identical localizations in the meiotic spindle, pole proteins have separable functions that impact pole stability in unique ways. Rather than all pole proteins interacting as a cohesive, cross-linked network around microtubule minus ends, it seems more plausible that different groups of proteins work in parallel to form and stabilize acentrosomal poles. These degron-based depletions with essential pole proteins have allowed for further mechanistic understanding of how each pole protein is contributing to spindle stability. There remain numerous pole proteins that could be subjected to rapid depletion via the AID system; the ability to probe the roles of essential proteins in both spindle assembly and maintenance is vital to building a more robust model of how meiotic spindles are able to achieve bipolarity in the absence of centrosomes to faithfully congress and segregate chromosomes.

## MATERIALS AND METHODS

### *C. elegans* strains used

| **<u>Name</u>** | **<u>Description</u>** | **<u>Genotype</u>** |
| --- | --- | --- |
| CA1215 | *dhc-1::degron::GFP*<br>*P<sub>sun-1</sub>::TIR1::mRuby* | *dhc-1(ie28[dhc-1::degron::GFP]) I; ieSi38 IV* |
| PHX3914 | *dhc-1::degron::GFP*<br>*P<sub>sun-1</sub>::TIR1::mRuby*<br>*GFP::H2B*<br>*GFP::tubulin*<br>*Δbmk-1* | *dhc-1(ie28[dhc-1::degron::GFP]) I; unc-119(ed3) III; ruls32[unc-119(+)*<br>*pie-1::gfp::h2b] III; ruls57[unc-119(+)*<br>*pie-1::GFP::tubulin]; ieSi38 IV; bmk-1(syb3914) V* |
| SMW22 | *P<sub>sun-1</sub>::TIR1::mRuby*<br>*mCherry::tubulin* | *unc-119(ed3) III; wels21 [pJA138 (pie-1::mCherry::tubulin::pie-1)]; ieSi38 IV* |
| SMW31 | *P<sub>sun-1</sub>::TIR1::mRuby*<br>*GFP::H2B*<br>*GFP::tubulin* | *unc-119(ed3) III; ruls32[unc-119(+)*<br>*pie-1::gfp::h2b] III; ruls57[unc-119(+)*<br>*pie-1::GFP::tubulin]; ieSi38 IV* |
| SMW46 | *dhc-1::degron::GFP*<br>*P<sub>sun-1</sub>::TIR1::mRuby*<br>*GFP::H2B*<br>*GFP::tubulin* | *dhc-1(ie28[dhc-1::degron::GFP]) I; unc-119(ed3) III; ruls32[unc-119(+)*<br>*pie-1::gfp::h2b] III; ruls57[unc-119(+)*<br>*pie-1::GFP::tubulin]; ieSi38 IV* |
| SMW47 | *dhc-1::degron::GFP*<br>*P<sub>sun-1</sub>::TIR1::mRuby*<br>*bmk-1(ok391)* | *dhc-1(ie28[dhc-1::degron::GFP]) I; ieSi38 IV; bmk-1(ok391) V* |
| SMW48 | *dhc-1::degron::GFP*<br>*P<sub>sun-1</sub>::TIR1::mRuby*<br>*mCherry::tubulin* | *dhc-1(ie28[dhc-1::degron::GFP]) I; wels21 [pJA138 (pie-1::mCherry::tub::pie-1)]; ieSi38 IV* |
| SV1005 | *bmk-1(ok391)* | *bmk-1(ok391) V* |

### Generation of *C. elegans* strains

CA1215 was generated via CRISPR/Cas9 editing of the endogenous *dhc-1* locus (Zhang et al., 2015).

PHX3914 was generated via CRISPR/Cas9 editing of the endogenous *bmk-1* locus by SunyBiotech.

SV1005 was a knockout strain generated by the *C. elegans* Deletion Mutant Consortium SMW46-SMW48 were generated via crossing of males and hermaphrodites of two strains and screening multiple generations of progeny to ensure homozygosity of all desired traits.

<u>SMW46:</u> Crossed males of SMW31 with CA1215 hermaphrodites

<u>SMW47:</u> Crossed males of CA1215 with SV1005 hermaphrodites

<u>SMW48:</u> Crossed males of CA1215 with SMW22 hermaphrodites

### RNAi Feeding

From a RNAi library (Kamath et al., 2003), individual RNAi clones were picked and grown overnight at 37°C in LB with 100μg/mL ampicillin. Overnight cultures were spun down, resuspended, and plated on nematode growth medium (NGM) plates containing 100μg/mL ampicillin and 1mM IPTG. Plates were dried in a dark space overnight at 22°C; at the same time, worms were synchronized for experimentation via bleaching of gravid adults, collecting remaining embryos, and plating on foodless plates to hatch overnight. The next day, newly hatched L1 worms were transferred to appropriate RNAi plates and grown to adulthood at 15°C for 5-6 days. In experiments utilizing *lin-5(RNAi)*, L1 worms were plated on EV (empty vector) control RNAi plates for 3 days, then transferred to *lin-5(RNAi)* plates 72 hours prior to fixation.

### Immunofluorescence and antibodies

Adult worms, grown on either EV (empty vector) control RNAi or experimental RNAi, were picked into a 10μL drop of Meiosis Media (Laband et al., 2018) in the center of a Poly-L-Lysine-coated glass slide. Worms were dissected, then fixed via freeze cracking and plunging into −20°C MeOH as described in (Oegema et al., 2001). Embryos were fixed for 40-45 minutes, rehydrated in PBS, and blocked in AbDil (PBS with 4% BSA, 0.1% Triton-X-100, 0.02% NaN_3_) overnight at 4°C. Primary antibodies were diluted in AbDil and incubated with sample overnight at 4°C. The following day, samples were moved to room temperature and rinsed three times in PBST (PBS with 0.1% Triton-X-100), then incubated with secondary antibodies (diluted in PBST) for 2 hours. Next, samples were washed three times in PBST again, and incubated with mouse anti-α-Tubulin-FITC (diluted in PBST) for 2 hours. Again, samples were washed three times in PBST, then incubated with Hoëscht (1:1000 in PBST) for 15 minutes. Finally, samples were washed two times in PBST, mounted in 0.5% *p*-phenylenediamine, 20mM Tris-Cl, pH 8.8, 90% glycerol, then sealed with nail polish and stored at 4°C.

Primary antibodies used in this study: rabbit-α-ASPM-1 (1:5000, gift from Arshad Desai), rabbit-α-BMK-1 (1:250, gift from Jill Schumacher), mouse-α-Tubulin-FITC (1:500, DM1α, Sigma), mouse-α-GFP (1:250, 3E6, Invitrogen), mouse-α-SUMO (1:500, gift from Federico Pelisch), and rabbit-α-SPD-1 (Mullen and Wignall, 2017). All rabbit and mouse Alexa-fluor secondary antibodies (Invitrogen) were used at 1:500.

### *Ex utero* live imaging

Fifteen adult worms, grown on either control RNAi or experimental RNAi, were picked into a 10μL drop of Meiosis Media in the center of a custom-made apparatus for live imaging (Laband et al., 2018; Divekar et al., 2021b). All worms were quickly dissected, and an eyelash pick was used to push remaining worm bodies to the outside of the drop, leaving only the embryos in the center to avoid disruption from worm movement during the imaging process.

Vaseline was laid in a ring around the drop through a syringe, and a 18×18mm #1 coverslip was laid on top of the Vaseline ring, sealing the drop. This sealed slide was moved immediately to the Spinning Disk stage and inverted, allowing embryos to float down to the surface of the coverslip and be subsequently imaged.

### Microscopy

All fixed imaging was performed on a DeltaVision Core deconvolution microscope with a 100x objective (NA = 1.4) (Applied Precision). This microscope is housed in the Northwestern University Biological Imaging Facility supported by the NU Office for Research. Image stacks were obtained at 0.2μm z-steps and deconvolved using SoftWoRx (Applied Precision). All immunofluorescence images in this study were deconvolved and displayed as full maximum intensity projections of data stacks encompassing the entire spindle structure (typically ∼4-6μm).

All live imaging was performed using a spinning disk confocal microscope with a 63x HC PL APO 1.40 NA objective lens. A spinning disk confocal unit (CSU-X1; Yokogawa Electric Corporation) attached to an inverted microscope (Leica DMI6000 SD) and a Spectral Applied Imaging laser merge ILE3030 and a back-thinned electron-multiplying charge-coupled device (EMCCD) camera (Photometrics Evolve 521 Delta) were used for image acquisition. The microscope and attached devices were controlled using Metamorph Image Series Environment software (Molecular Devices). Typically, ten to fifteen z-stacks at 1μm increments were taken every 15–30 seconds at room temperature. Images were processed using ImageJ; images are shown as maximum intensity projections of the entire spindle structure. The spinning disk microscope is housed in the Northwestern University Biological Imaging Facility supported by the NU Office for Research.

### Acute auxin treatment of *C. elegans*

Acute auxin treatments utilized in immunofluorescence experiments were performed as in (Divekar et al., 2021a). Briefly, a Meiosis Media solution containing 1mM auxin was prepared using a 200mM stock of auxin dissolved in 100% EtOH and kept on ice. Worms were picked into 10μL drops of Media+auxin, and then the slides were placed inside a custom-made humidity chamber to avoid evaporation of the drop. Following incubation for desired time (30-45 minutes), worms were immediately dissected and subjected to standard IF protocol described above.

For acute auxin treatments utilized in *ex utero* live imaging, a Meiosis Media solution containing 100μM auxin was made from a 200mM stock of auxin dissolved in 100% EtOH. Worms were picked into 10μL drops of Media+auxin, and dissected on a custom-made live imaging slide apparatus described above. Slides were moved quickly to the Spinning Disk, and acquisition was started as soon as an embryo could be found in order to begin imaging prior to full protein depletion. More detailed auxin treatment protocols are described in (Divekar et al., 2021b).

### Ethanol Fixation for Germline Counting

SMW46 worms were subjected to EV Control RNAi, utilizing methods described above, and were grown to adulthood over 5 days. Four hours prior to fixation, worms were either transferred to EV Control RNAi (with 1mM auxin) or left on their original RNAi plates. For each biological replicate, ∼40 adults were picked off their respective plates into a 10μL drop of M9 on a standard glass slide, and a small piece of Whatman paper was used to absorb excess M9, with care being used to avoid pulling worms onto filter paper. Once worms had formed a tight cluster with little residual M9, a 10μL drop of 100% EtOH was quickly pipetted onto the worms. Within seconds, worms became rigid and straight; once the first EtOH drop had dried, another 10μL drop was applied and allowed to completely dry. After the third drop was added and dried, a 10μL drop of 50% Vectashield Mounting Media (Vector Laboratories H-1000) and 50% M9 was placed onto the worms, and a 18×18mm coverslip was gently placed on top. Excess media was aspirated away, slides were sealed with nail polish, and stored at 4°C (typically imaged within a week of fixation).

### Data analysis

**Figure 1D:** Quantifications were made by viewing whole Dynein AID worms expressing GFP::tubulin and GFP::histone and observing three positions in each worm (−1 Oocyte, Spermatheca, +1 Embryo) in both gonad arms. When spindle morphology was clear enough to confidently categorize, embryos (already sorted based on location) were then counted into one of four categories (Microtubule Cage, Unfocused, Focused, Anaphase). Number of embryos counted across all slides (n) are placed above each respective location and condition. Each bar comprises embryo counts from at least 4 biological replicates.

**Figure 1E and 1F:** Categorization of acentrosomal poles into either focused or unfocused/splayed was done by eye, looking at both microtubule and ASPM-1 channels. In any case where microtubule bundles were clearly separated from one another at an acentrosomal pole, with multiple ASPM-1 foci, that spindle was considered unfocused/splayed. Quantifications of acentrosomal pole splaying were made by categorizing multiple fixed images of embryos; numbers of embryos counted across all slides (n) are placed above each respective condition. Each bar comprises embryos imaged across 3 biological replicates.

**Figure 2A and 2B:** Categorization of acentrosomal poles into either focused or unfocused/splayed was done by eye, looking at both microtubule and ASPM-1 channels. In any case where microtubule bundles were clearly separated from one another at an acentrosomal pole, with multiple ASPM-1 foci, that spindle was considered unfocused/splayed. Quantifications of acentrosomal pole splaying were made by categorizing multiple fixed images of embryos; numbers of embryos counted across all slides (n) are placed above each respective condition. Each bar comprises embryos imaged across at least 3 biological replicates.

**Figure 2 – figure supplement 1A and 1B:** Utilizing FIJI, a rectangular ROI (12μm or 14μm wide by 4μm tall) was used on all images of a single condition to measure the average pixel intensity of ASPM-1 across the whole width of the ROI via the Plot Profile Tool. These values were then averaged from all images in that condition to produce a single value for ASPM-1 intensity at each pixel across the standardized 12μm/14μm length. This was plotted as a solid line for each condition, with shaded areas above and below the line representing the SEM. All images were taken with the same exposures for all channels. Spindles were sum projected over 20 slices (0.2μm step size) for each image prior to measurement. Each condition was averaged using images from at least 3 biological replicates. Spindle length measurements were done in Imaris. Poles were projected into 3D volumes using the Surfaces Tool (based off of fluorescence intensity), and the center of each volume was determined. The exact micron distance between the designated center of the two volumes was measured, and this distance was averaged across images of the same condition. Quantifications were arranged as boxplots, and statistical significance was determined via a two-tailed t test. Each condition was averaged using images from at least 3 biological replicates.

**Figure 3A:** Quantifications were made by viewing whole Dynein AID worms expressing GFP::tubulin and GFP::histone and observing all +1 embryos. When spindle morphology was clear enough to confidently categorize, embryos were then counted as either monopolar (a single structure with chromosomes fanned away from center) or undergoing breakdown (noticeable separation of microtubules and chromosomes). Number of embryos counted across all slides (n) are placed above each respective location and condition. Each bar comprises embryo counts from 3 biological replicates.

**Figure 3 - figure supplement 1A:** Quantifications were made by viewing whole Dynein AID worms expressing GFP::tubulin and GFP::histone and observing two positions in each worm (Spermatheca and +1 Embryo) in both gonad arms. When spindle morphology was clear enough to confidently categorize, embryos (already sorted based on location) were then counted into one of three categories (Unfocused, Focused, Anaphase). Number of embryos counted across all slides (n) are placed above each respective location and condition. Each bar comprises embryo counts from at least 3 biological replicates.

**Figure 4 - figure supplement 1:** Timepoints from live imaging videos were analyzed utilizing Imaris. Chromosomes were rendered into 3D surfaces using the “Surfaces” tool, and the center of each volume was determined. For each chromosome that was within the z-stack throughout the entirety of the timelapse, measurements of the distance between the center of segregating chromosomes were taken. These distances were averaged at each 15 second timepoint, and each timelapse was standardized to the same starting point (onset of Anaphase A) to allow for further averaging between separate videos. The average segregation distance across each condition was plotted (number of videos in each condition represented by n), and shaded area represents SEM of each average.

**Figure 6B and 6C:** Classification of monopolar spindles into one of three categories (Breakdown, Mini Anaphases, No Segregation) was done by eye, looking at DNA, microtubule, and ASPM-1 channels. In any case where some number of bivalents were clearly separated from the monopolar spindle, yet a distinct ASPM-1 monopole remained with some number of attached bivalents, was classified as “Breakdown”. Whenever all six bivalents could be seen dispersed in the cytoplasm with no remaining ASPM-1 monopole, and appeared to have some segregation between chromosomes, that embryo was considered “Mini Anaphases”. Any oocytes containing dispersed chromosomes that were observed without any indication of segregation or an anaphase spindle were classified as “No Segregation”. Numbers of embryos counted across all slides (n) are placed above each respective condition. Each bar comprises embryos imaged across 5 biological replicates.

**Table 1.**
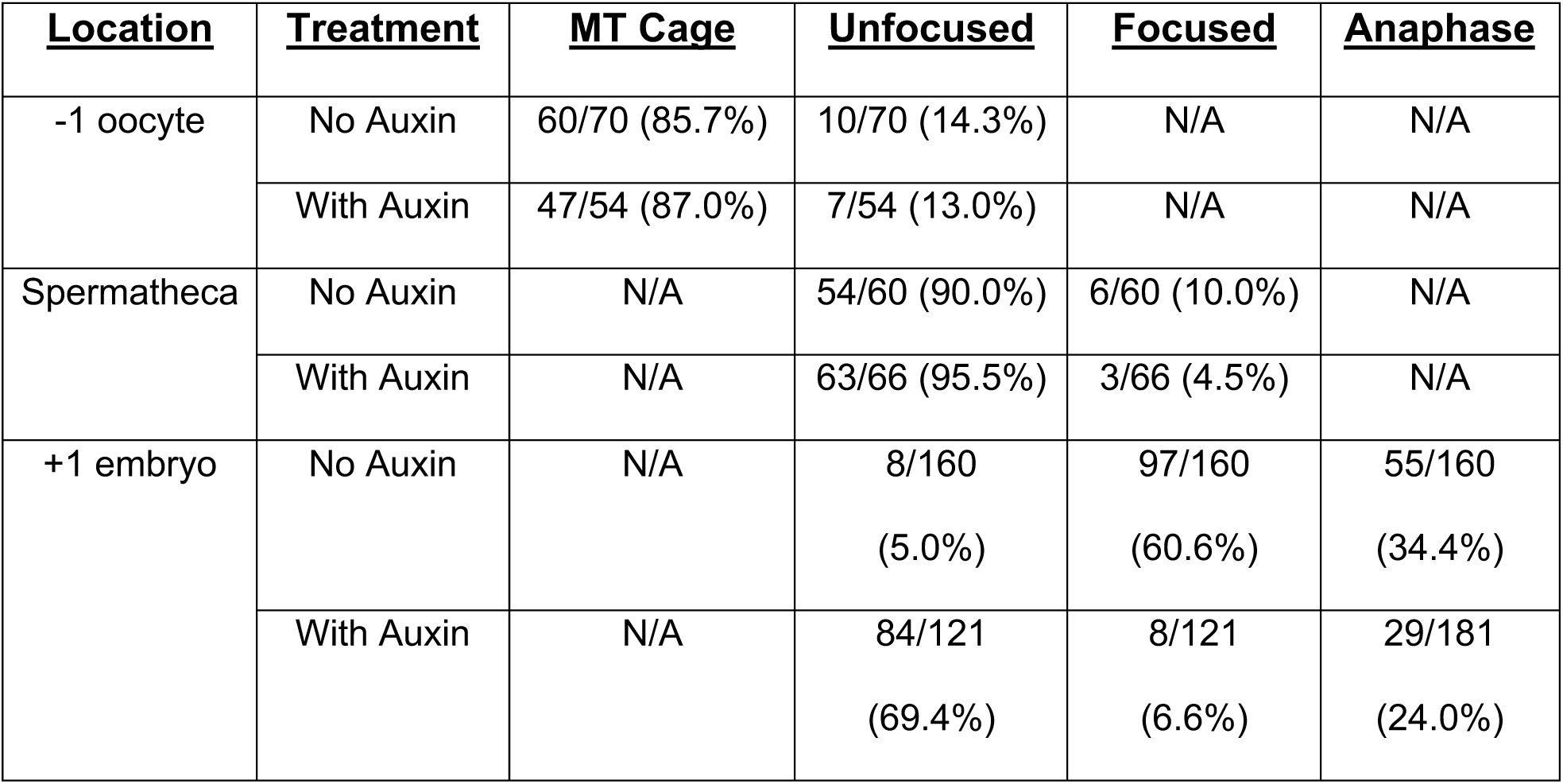
(corresponds to Figure 1D)

| <u>Location</u> | <u>Treatment</u> | <u>MT Cage</u> | <u>Unfocused</u> | <u>Focused</u> | <u>Anaphase</u> |
| --- | --- | --- | --- | --- | --- |
| -1 oocyte | No Auxin | 60/70 (85.7%) | 10/70 (14.3%) | N/A | N/A |
|  | With Auxin | 47/54 (87.0%) | 7/54 (13.0%) | N/A | N/A |
| Spermatheca | No Auxin | N/A | 54/60 (90.0%) | 6/60 (10.0%) | N/A |
|  | With Auxin | N/A | 63/66 (95.5%) | 3/66 (4.5%) | N/A |
| +1 embryo | No Auxin | N/A | 8/160<br>(5.0%) | 97/160<br>(60.6%) | 55/160<br>(34.4%) |
|  | With Auxin | N/A | 84/121<br>(69.4%) | 8/121<br>(6.6%) | 29/181<br>(24.0%) |

**Table 2.** (corresponds to Figure 1F)

| <u>Treatment</u> | <u>Focused Poles</u> | <u>Unfocused Poles</u> | <u>Anaphases</u> |
| --- | --- | --- | --- |
| No Auxin | 35/60 (58.3%) | 3/60 (5.0%) | 22/60 (36.7%) |
| With Auxin | 3/54 (5.6%) | 39/54 (72.2%) | 12/54 (22.2%) |

**Table 3.**
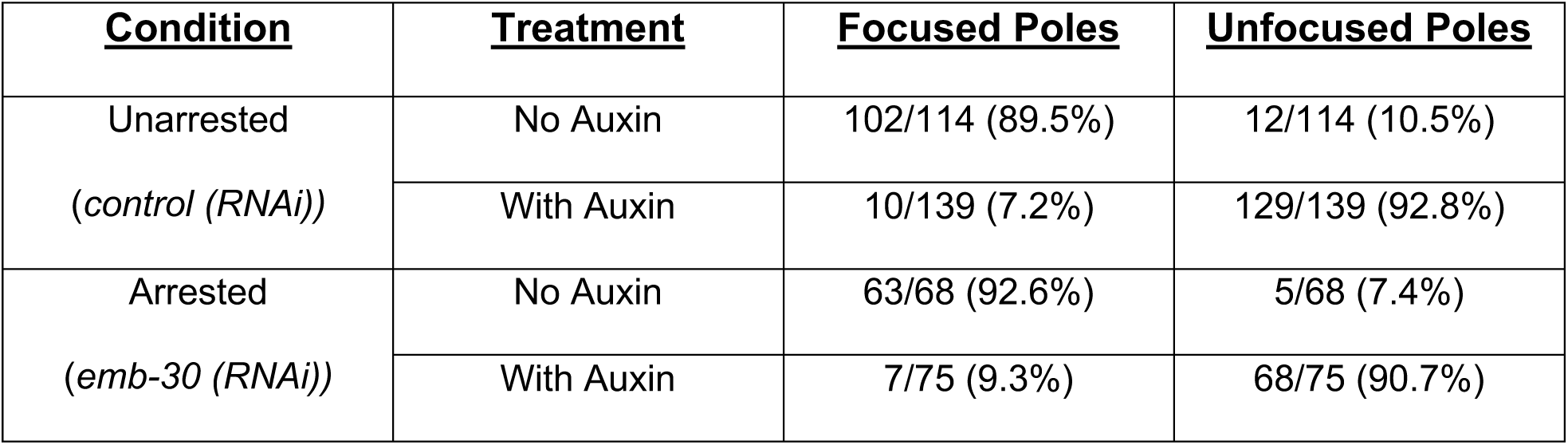
(corresponds to Figure 2B)

**Table 4.** (corresponds to Figure 3A)

| **<u>Condition</u>** | **<u>Treatment</u>** | **<u>Monopolar</u>** | **<u>Breakdown</u>** |
| --- | --- | --- | --- |
| *klp-18 (RNAi)* | No Auxin | 119/119 (100.0%) | 0/119 (0.0%) |
|  | With Auxin | 9/125 (7.2%) | 116/125 (92.8%) |

**Table 5.** (corresponds to Figure 3C)

| **<u>Condition</u>** | **<u>Intact Monopole</u>** | **<u>Partial Breakdown</u>** | **<u>Full Breakdown</u>** |
| --- | --- | --- | --- |
| *aspm-1 (RNAi);*<br>*klp-18 (RNAi)* | 18/61 (29.5%) | 29/61 (47.5%) | 14/61 (23.0%) |

**Table 6.** (corresponds to Figure 3 – figure supplement 1A)

| **<u>Location</u>** | **<u>Condition</u>** | **<u>Focused Poles</u>** | **<u>Unfocused Poles</u>** | **<u>Anaphases</u>** |
| --- | --- | --- | --- | --- |
| Spermatheca | *control (RNAi)* | 5/71 (7.0%) | 66/71 (93.0%) | N/A |
|  | *lin-5 (RNAi)* | 4/83 (4.8%) | 79/83 (95.2%) | N/A |
| +1 embryo | *control (RNAi)* | 72/115 (62.6%) | 9/115 (7.8%) | 34/115 (29.6%) |
|  | *lin-5 (RNAi)* | 10/188 (5.3%) | 120/188 (63.8%) | 58/188 (30.9%) |

**Table 7.** (corresponds to Figure 3 – figure supplement 1B)

| **<u>Condition</u>** | **<u>Intact Monopole</u>** | **<u>Partial Breakdown</u>** | **<u>Full Breakdown</u>** |
| --- | --- | --- | --- |
| *lin-5 (RNAi);*<br>*klp-18 (RNAi)* | 14/73 (19.2%) | 44/73 (60.3%) | 15/73 (20.5%) |

**Table 8.** (corresponds to Figure 6C)

| <u>Condition</u> | <u>Treatment</u> | <u>Breakdown</u> | <u>Mini Anaphases</u> | <u>No Segregation</u> |
| --- | --- | --- | --- | --- |
| *klp-18 (RNAi)* | *bmk-1 (WT)* | 82/130 (63.1%) | 43/130 (33.1%) | 5/130 (3.8%) |
|  | *bmk-1 (ok391)* | 111/164 (67.7%) | 0/164 (0.0%) | 53/164 (32.3%) |

## ACKNOWLEDGMENTS

We thank all members of the Wignall lab for support and discussion, and Emily Czajkowski, Nikita Divekar, Hannah Horton, Juhi Narula, and Ian Wolff for critical reading of the manuscript. Additionally, we specifically thank graduate student Karlin Compton for their help in piloting certain experimental conditions. We thank SunyBiotech for their service in generating PHX3914 for live imaging experiments. Finally, we thank the Dernburg Lab for providing CA1215, the original *dhc-1::degron::GFP* worm strain, and Arshad Desai, Federico Pelisch, and Jill Schumacher for antibodies. Some strains were provided by the *Caenorhabditis* Genetics Center (CGC), which is funded by NIH Office of Research Infrastructure Programs (P40 OD010440). This work was funded by NIH R01GM124354 (to SMW) and by NIH/NCI training grant T32 CA009560 (to GCM).

---

**Figure 2 – figure supplement 1.**
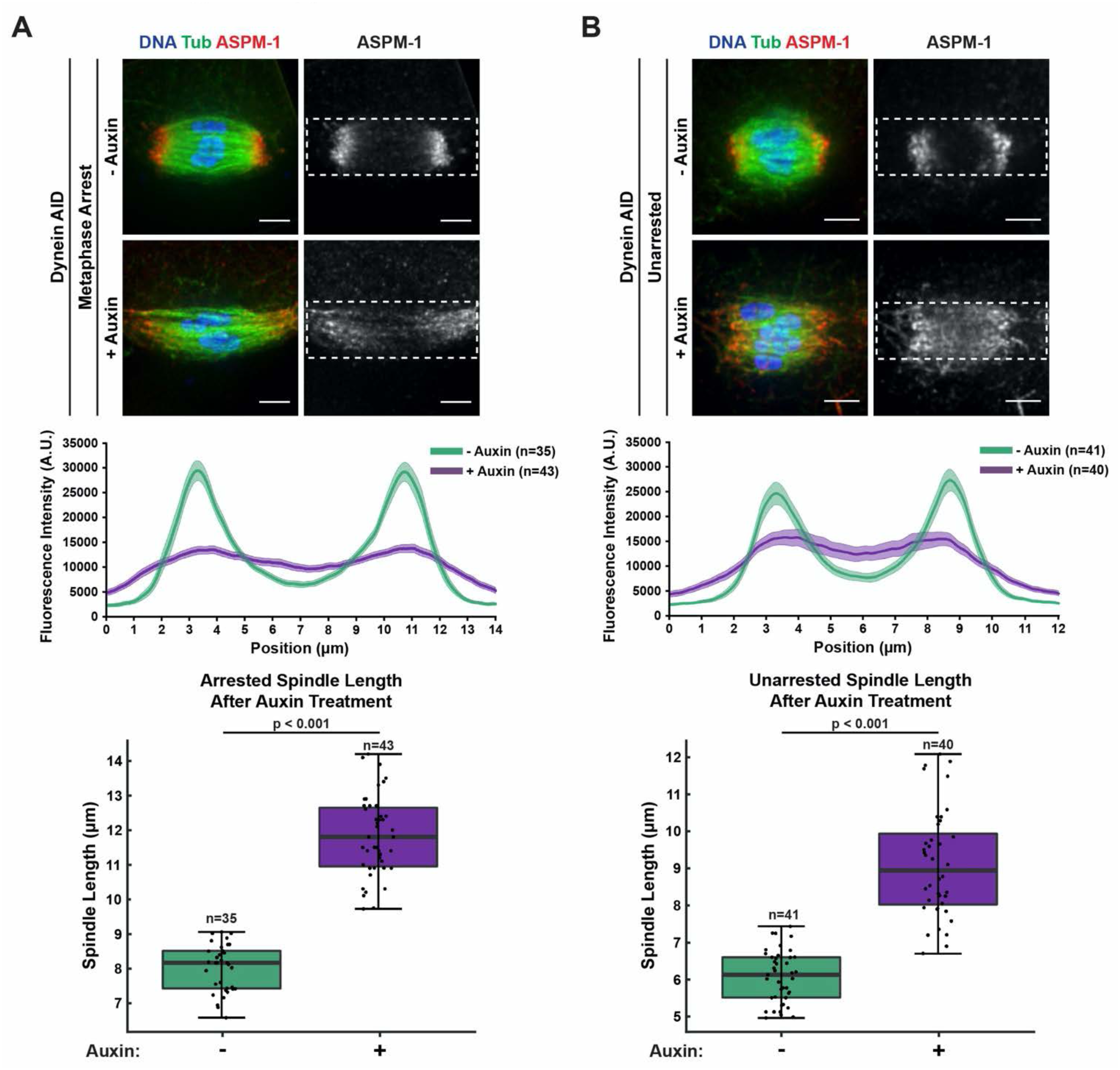
Acute dynein depletion leads to dispersion of ASPM-1 across the spindle and increased pole-to-pole spindle length. Representative IF images showing the phenotype of acute dynein depletion from metaphase-arrested *(emb-30(RNAi))* spindles (A) or control spindles (B) in Dynein AID worms; shown are microtubules (green), DNA (blue), and ASPM-1 (red). Dashed rectangles represent ROI used for linescan measurements (4μm tall by 12 or 14μm wide). Quantifications of fluorescence intensity across the length of the spindle demonstrate that dynein depletion leads to considerably more uniform and dispersed ASPM-1 localization. All images used for linescan analysis were also measured for pole-to-pole spindle length, and these measurements were plotted in corresponding boxplots. Upon addition of auxin, the spindle is significantly longer. All scale bars = 2.5μm. Statistical significance between means of spindle length was determined via a two-tailed t test for unarrested and arrested conditions, with a p value of p<2.2 x 10^-16^ for both.

**Figure 2 – figure supplement 2.**
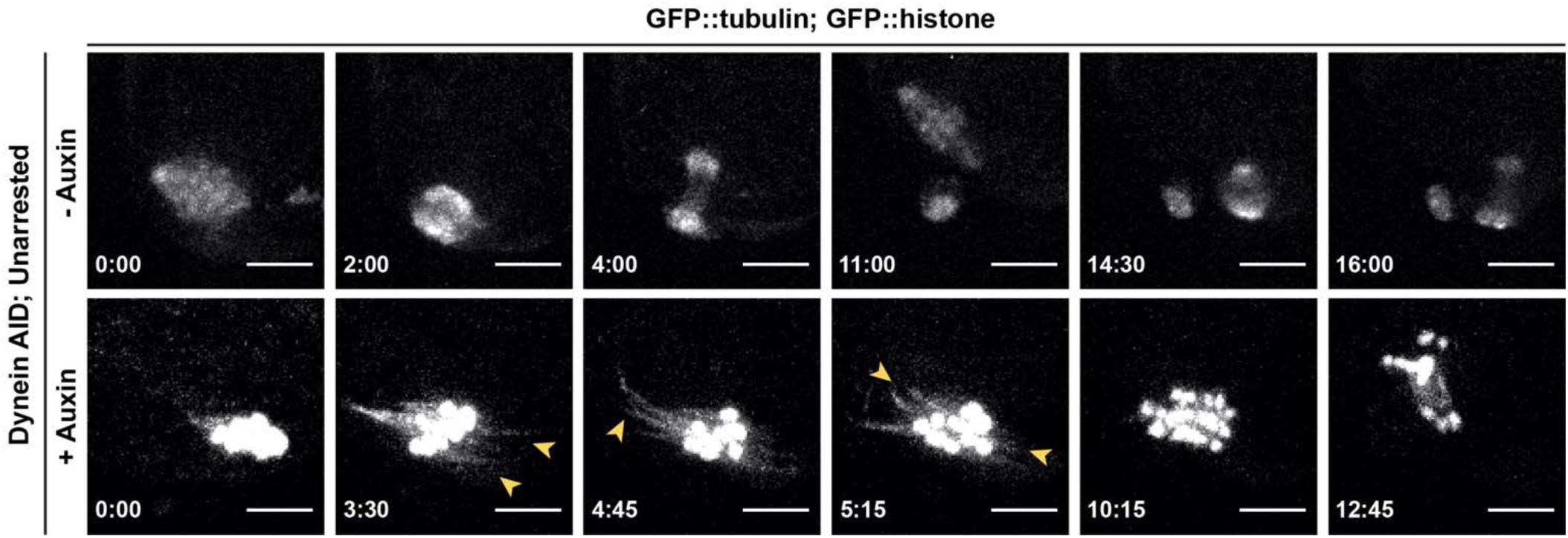
Phenotypes seen in dynein depletion are not an artifact of metaphase arrest. Repeat of *ex utero* live imaging in Figure 2D, but with unarrested embryos. Splaying of spindle poles (arrowheads) is identical to splaying observed in metaphase-arrested spindles (Figure 2C**, 2D**). We also observed spindle rotation defects in anaphase, consistent with prior studies. Time elapsed shown in (min):(sec). Scale bars = 5µm.

**Figure 3 – figure supplement 1.**
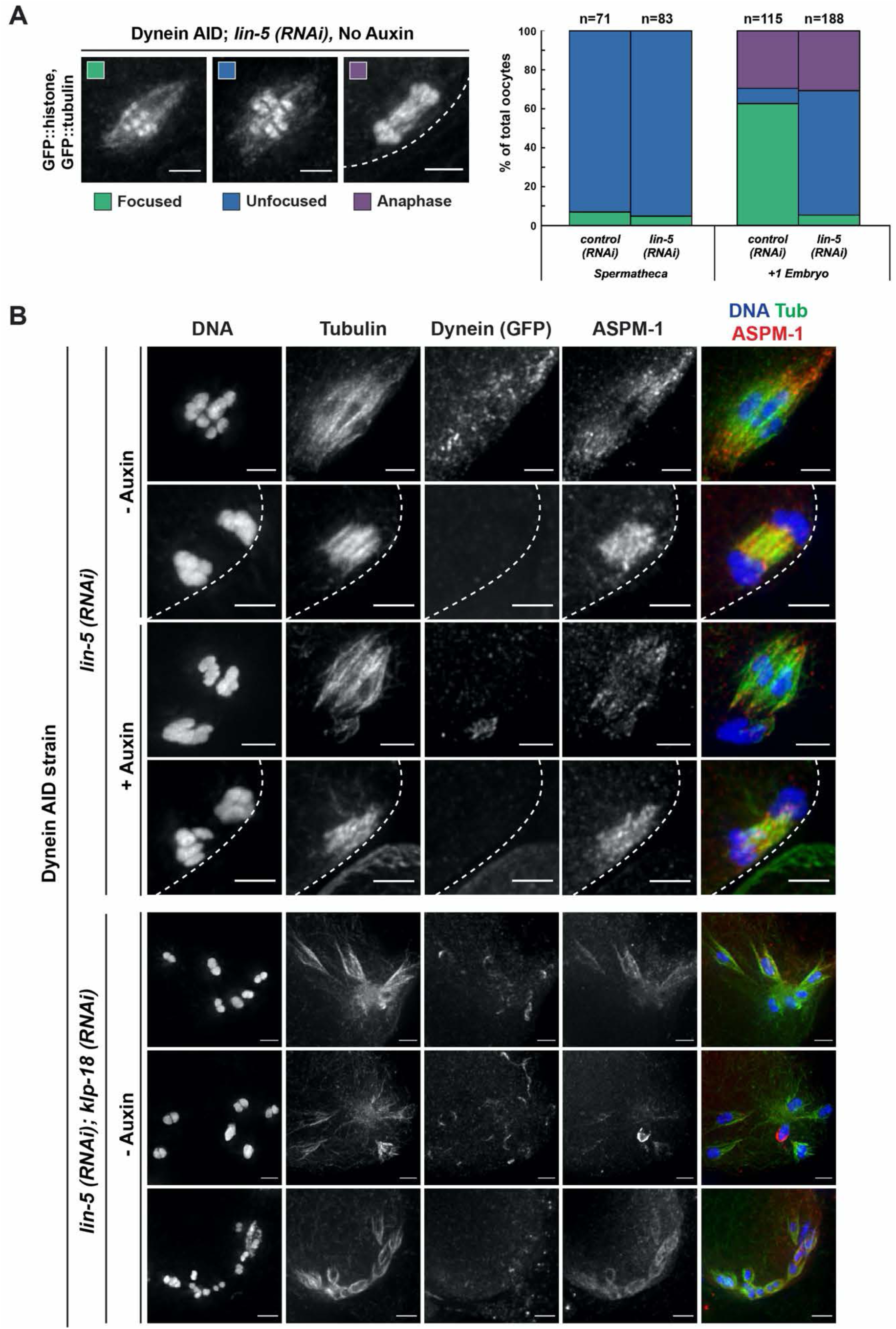
Meiotic spindles following LIN-5 depletion are morphologically similar to Dynein AID conditions. A) Representative images of spindles counted in germline imaging from the Dynein AID strain expressing GFP::tubulin and GFP::histone. In *lin-5(RNAi)* conditions, splaying of acentrosomal poles can be observed in the +1 position. We also observed improper spindle rotation during anaphase, consistent with prior studies. Quantifications demonstrated similar ratios of unfocused poles as those seen in Dynein AID germline counting. Cortex represented by dashed line. B) IF imaging of embryos from Dynein AID worms in *lin-5* RNAi conditions; shown are tubulin (green), DNA (blue), ASPM-1 (red) and dynein (not shown in merge). ASPM-1 labeling further supports splaying of acentrosomal poles and improper spindle rotation during anaphase can also be seen. Concurrent acute dynein depletion (rows 3, 4) does not seem to exacerbate the spindle defects observed in *lin-5(RNAi)* alone. Double RNAi depletions with *klp-18(RNAi)* cause similar breakdown phenotypes to Dynein AID depletions on monopoles (rows 5, 6, 7). Cortex represented by dashed line. All scale bars = 2.5μm.

**Figure 4 – figure supplement 1.**
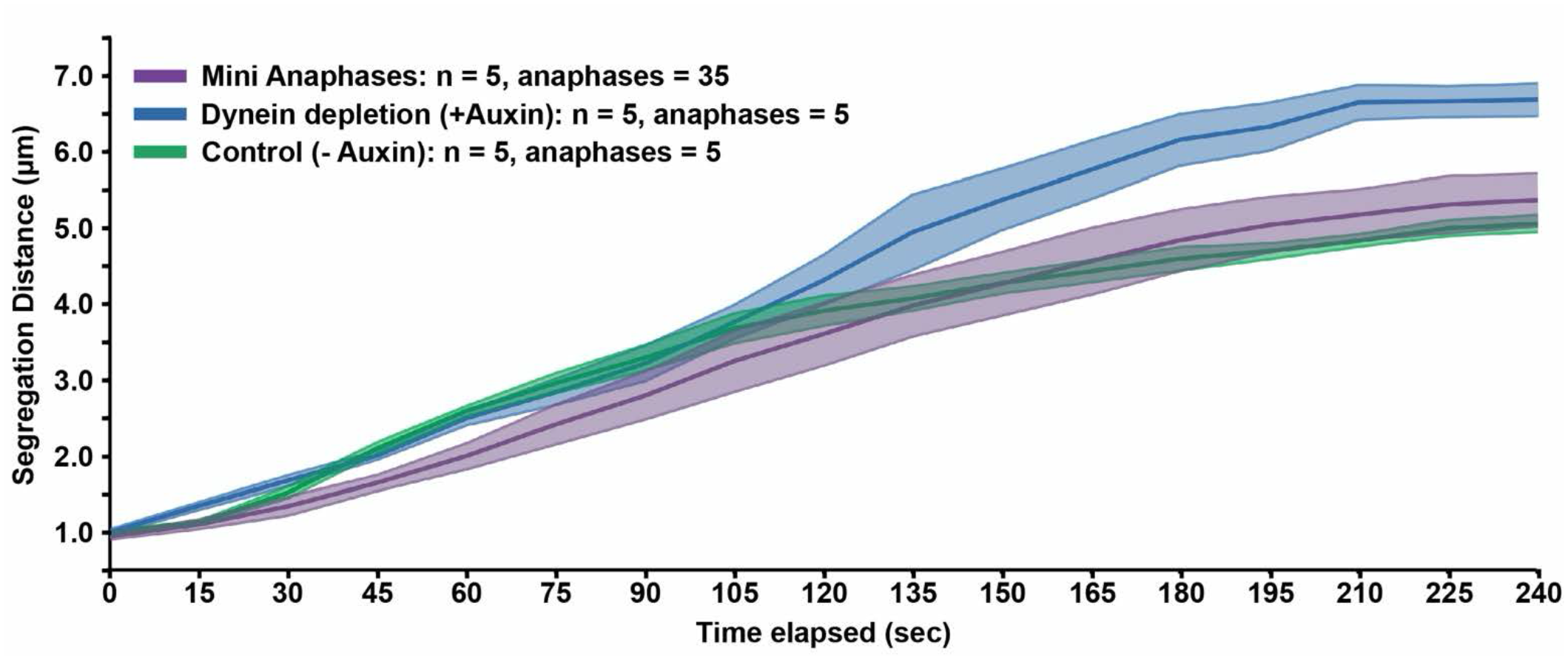
Miniature anaphases segregate chromosomes at distances and rates comparable to normal anaphase spindles. Quantification of timelapses from control anaphases (the Dynein AID strain without auxin), anaphases in the absence of dynein (the Dynein AID strain with auxin), and miniature anaphases. Each timelapse was synchronized to initiation of anaphase A, and the distance between the centers of each chromosome was measured at each 15 second timepoint. For each condition, the number of timelapses used is represented by n, and the shaded area around each average line represents SEM for each condition. Note that for the mini anaphase condition, though we used 5 movies, we measured multiple mini anaphases in each (total anaphases = 35). Miniature anaphases do not exhibit significant differences in segregation rates or distances to wild type anaphase spindles, but do not reach distances of dynein depletions alone.

**Figure 5 – figure supplement 1.**
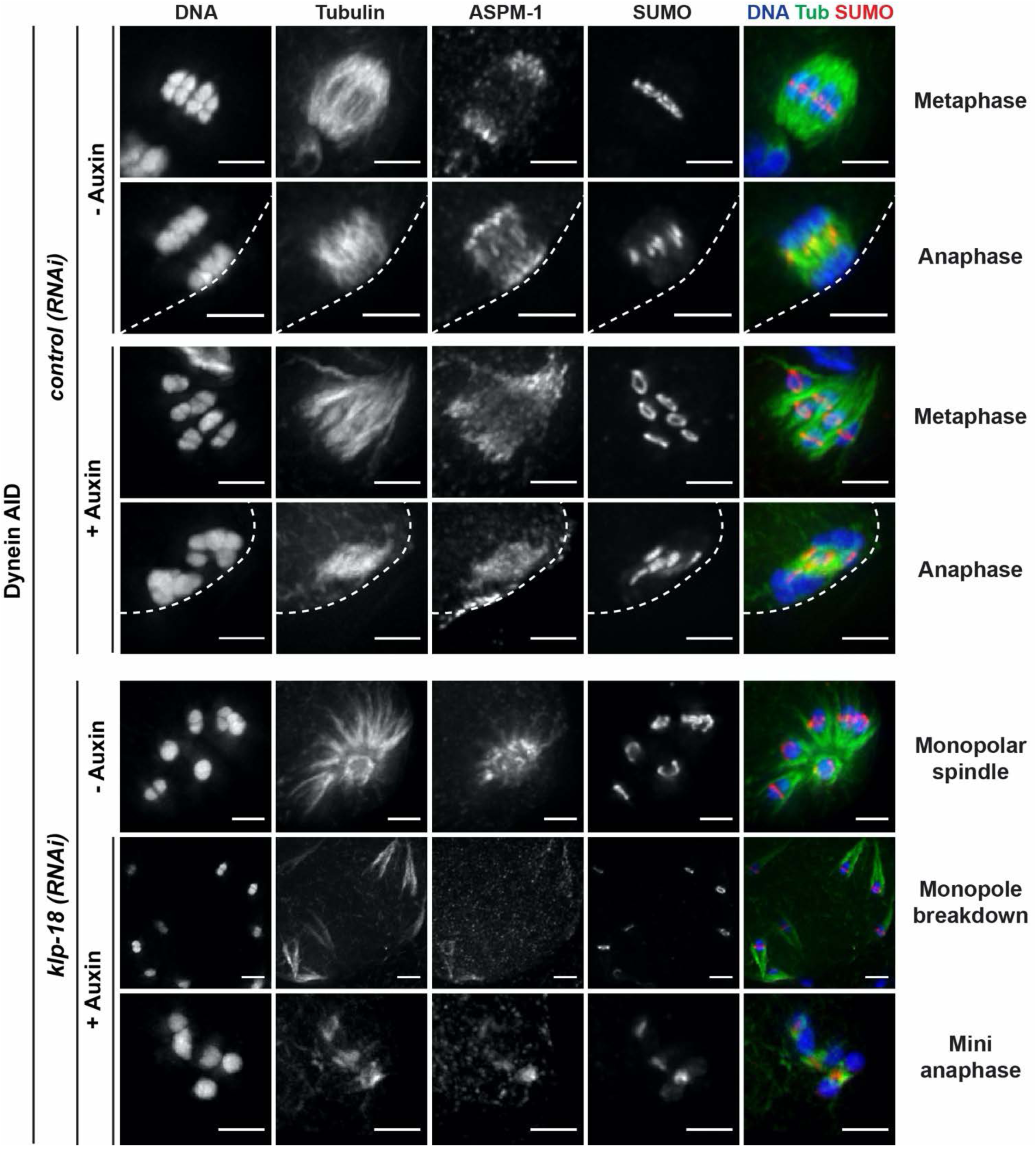
SUMO is localized normally in all dynein depletion conditions. IF imaging of control and *klp-18(RNAi)* conditions; shown are tubulin (green), DNA (blue), SUMO (red) and ASPM-1 (not shown in merge). SUMO is localized to a ring complex that forms around chromosomes and then is released from chromosomes in anaphase in the presence or absence of dynein (rows 1-4). SUMO remains localized to the ring complex before and after monopole breakdown (rows 5, 6), and can clearly be seen localized between separating chromosomes in miniature anaphases (row 7). Cortex is represented by dashed line. Scale bars = 2.5µm.

**Figure 6 – figure supplement 1.**
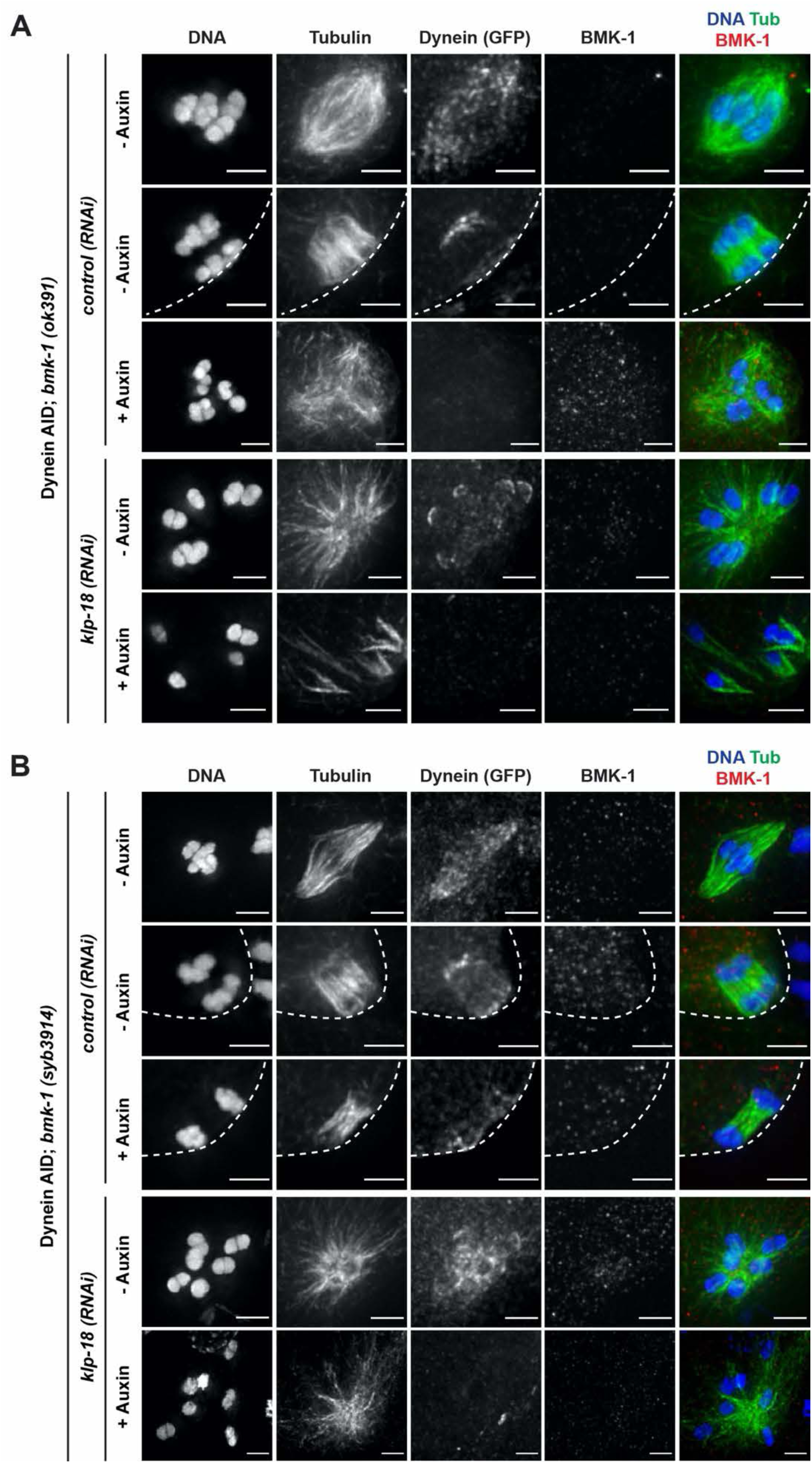
IF imaging validation of BMK-1 deletion in worm strains. IF imaging of embryos containing either the *bmk-1(ok391)* deletion allele (A) or the *bmk-1(syb3914)* deletion allele (B); shown are tubulin (green), DNA (blue), BMK-1 (red) and dynein (not shown in merge). No BMK-1 localization can be observed in either mutant in *control(RNAi)* or *klp-18(RNAi)* conditions in the presence or absence of auxin. Cortex represented by dashed line. Scale bars = 2.5μm.

## SUPPLEMENTAL VIDEOS

**Video 1. Localization of dynein during oocyte meiosis.**

Shows an embryo expressing mCherry::tubulin, dissected into a control Meiosis Media solution. DHC-1::degron::GFP localization can be seen at spindle poles during metaphase, alongside faint localization to kinetochores. Localization is consistent across all embryos filmed (n=2). Time elapsed shown in (min):(sec). Scale bar = 5μm.

**Video 2. Dynein localizes to spindle poles in metaphase-arrested spindles.**

Shows a metaphase-arrested embryo expressing mCherry::tubulin (top), dissected into a control Meiosis Media solution. DHC-1::degron::GFP (bottom) is clearly localized to spindle poles, and faintly seen at kinetochores. Corresponds to Figure 2C. Localization is consistent across all embryos filmed (n=3). Time elapsed shown in (min):(sec). Scale bar = 5μm.

**Video 3. Auxin treatment rapidly depletes dynein and unfocuses poles.**

Shows a metaphase-arrested embryo expressing mCherry::tubulin (top), dissected into Meiosis Media containing 100μM auxin to deplete dynein. Once dissected into auxin solution, rapid depletion of DHC-1::degron::GFP (bottom) is evident within 3 minutes, at which point spindle poles begin to unfocus and splay apart. Corresponds to Figure 2C. Phenotypes are consistent across all embryos filmed (n=4). Time elapsed shown in (min):(sec). Scale bar = 5μm.

**Video 4. Metaphase-arrested spindles remain stable and maintain chromosome alignment.**

Shows a metaphase-arrested embryo expressing GFP::tubulin and GFP::histone, dissected into a control Meiosis Media solution. No major changes in spindle length or shape occur and chromosomes are stably aligned in the spindle center. Corresponds to Figure 2D. Phenotypes are consistent across all embryos filmed (n=5). Time elapsed shown in (min):(sec). Scale bar = 5μm.

**Video 5. Auxin treatment unfocuses poles, but spindle and chromosomes largely retain alignment along a single axis.**

Shows a metaphase-arrested embryo expressing GFP::tubulin and GFP::histone, dissected into Meiosis Media containing 100μM auxin to deplete dynein. Upon dissection into auxin solution, rapid and dynamic splaying of acentrosomal poles occurs alongside a notable increase in spindle length, but microtubule bundles and chromosomes stay somewhat aligned along a single axis. Corresponds to Figure 2D. Phenotypes are consistent across all embryos filmed (n=6). Time elapsed shown in (min):(sec). Scale bar = 5μm.

**Video 6. Normal embryos undergo two rounds of bipolar spindle formation and anaphase segregation.**

Shows an unarrested embryo expressing GFP::tubulin and GFP::histone, dissected into a control Meiosis Media solution. After one round of segregation, a second bipolar spindle is formed, rotates perpendicular to the cortex, and successfully extrudes a second polar body. Corresponds to Figure 2 – figure supplement 2. Phenotypes are consistent across all embryos filmed (n=5). Time elapsed shown in (min):(sec). Scale bar = 5μm.

**Video 7. Splaying of acentrosomal poles occurs in unarrested embryos.**

Shows an unarrested embryo expressing GFP::tubulin and GFP::histone, dissected into Meiosis Media containing 100μM auxin to deplete dynein. A spindle tries to form in the absence of dynein, leading to dynamic and unfocused poles. Anaphase segregation still occurs, albeit with a failure in spindle rotation. Corresponds to Figure 2 – figure supplement 2. Phenotypes are consistent across all embryos filmed (n=7). Time elapsed shown in (min):(sec). Scale bar = 5μm.

**Video 8. Complete dissolution of monopoles occurs upon auxin addition.**

Shows an embryo expressing GFP::tubulin and GFP::histone following *klp-18(RNAi)*, dissected into Meiosis Media containing 100μM auxin to deplete dynein. Upon dissection into auxin solution, the monopole quickly dissolves, leading to ejection of individual chromosomes (with laterally-associated microtubule bundles) into the cytoplasm, leading to an anaphase-like segregation event. Corresponds to Figure 4A. Phenotypes are consistent across all embryos filmed (n=5). Time elapsed shown in (min):(sec). Scale bar = 5μm.

**Video 9. Dynamics of normal monopolar spindle.**

Shows an embryo expressing GFP::tubulin and GFP::histone following *klp-18(RNAi)*, dissected into a Meiosis Media control solution. Chromosomes extend outward towards microtubule plus ends forming a monopolar spindle during metaphase, but retract towards the minus ends during anaphase. Corresponds to Figure 4A. Phenotypes are consistent across all embryos filmed (n=6). Time elapsed shown in (min):(sec). Scale bar = 5μm.

**Video 10. Microtubule reorganization leads to anaphase-like segregation for individual chromosomes.**

Shows an embryo expressing GFP::tubulin and GFP::histone following *klp-18(RNAi)*, dissected into Meiosis Media containing 100μM auxin to deplete dynein. After breakdown of the monopolar spindle, reorganization of local microtubule bundles around individual chromosomes can be observed. Shortly after reorganization of these miniature spindles, an anaphase-like segregation occurs synchronously across all chromosomes. Corresponds to Figure 4B. Phenotypes are consistent across all embryos filmed (n=5). Time elapsed shown in (min):(sec). Scale bar = 5μm.

**Video 11. Monopolar spindle dynamics are normal in the absence of BMK-1.**

Shows a *bmk-1(syb3914)* embryo expressing GFP::tubulin and GFP::histone following *klp-18(RNAi)* dissected into a control Meiosis Media solution. Without functional BMK-1 or KLP-18, monopolar spindles can still form and monopolar anaphase still occurs. Phenotypes are consistent across all embryos filmed (n=4). Time elapsed shown in (min):(sec). Scale bar = 5μm.

**Video 12. Loss of BMK-1 function prevents the formation of miniature anaphases.** Shows a *bmk-1(syb3914)* embryo expressing GFP::tubulin and GFP::histone following *klp-18(RNAi)* dissected into Meiosis Media containing 100μM auxin to deplete dynein. Without functional BMK-1, formation of miniature bipolar anaphases do not occur after the breakdown of the monopolar spindle. Corresponds to Figure 6D. Phenotypes are consistent across all embryos filmed (n=3). Time elapsed shown in (min):(sec). Scale bar = 5μm.

